# Feedforward extraction of behaviorally significant information by neocortical columns

**DOI:** 10.1101/2025.04.20.649709

**Authors:** Oleg V. Favorov, Olcay Kursun

## Abstract

Neurons throughout the neocortex exhibit selective sensitivity to particular features of sensory input patterns. According to the prevailing views, cortical strategy is to choose features that exhibit predictable relationship to their spatial and/or temporal context. Such contextually predictable features likely make explicit the causal factors operating in the environment and thus they are likely to have perceptual/behavioral utility. The known details of functional architecture of cortical columns suggest that cortical extraction of such features is a modular nonlinear operation, in which the input layer, layer 4, performs initial nonlinear input transform generating proto-features, followed by their linear integration into output features by the basal dendrites of pyramidal cells in the upper layers. Tuning of pyramidal cells to contextually predictable features is guided by the contextual inputs their apical dendrites receive from other cortical columns via long-range horizontal or feedback connections. Our implementation of this strategy in a model of prototypical V1 cortical column, trained on natural images, reveals the presence of a limited number of contextually predictable orthogonal basis features in the image patterns appearing in the column’s receptive field. Upper-layer cells generate an overcomplete Hadamard-like representation of these basis features: i.e., each cell carries information about all basis features, but with each basis feature contributing either positively or negatively in the pattern unique to that cell. In tuning selectively to contextually predictable features, upper layers perform selective filtering of the information they receive from layer 4, emphasizing information about orderly aspects of the sensed environment and downplaying local, likely to be insignificant or distracting, information. Altogether, the upper-layer output preserves fine discrimination capabilities while acquiring novel higher-order categorization abilities to cluster together input patterns that are different but, in some way, environmentally related. We find that to be fully effective, our feature tuning operation requires collective participation of cells across 7 minicolumns, together making up a functionally defined 150μm diameter “mesocolumn.” Similarly to real V1 cortex, 80% of model upper-layer cells acquire complex-cell receptive field properties while 20% acquire simple-cell properties. Overall, the design of the model and its emergent properties are fully consistent with the known properties of cortical organization. Thus, in conclusion, our feature-extracting circuit might capture the core operation performed by cortical columns in their feedforward extraction of perceptually and behaviorally significant information.

## 1 INTRODUCTION

In artificial intelligence, to quote Ritter (2001), “A first and very important step in many pattern recognition and information processing tasks is the identification or construction of a reasonably small set of important features in which the essential information for the task is concentrated.” The concept of stimulus feature tuning is also fundamental to neuroscience. It is widely accepted that neurons throughout the cerebral cortex exhibit highly selective sensitivity to particular features of peripheral stimuli and such tuning defines neurons’ information representational identities (DiCarlo and Cox, 2007). Across cortical areas, neurons tune to stimulus features that vary greatly in their complexity. Starting from the primary sensory cortical areas and up each stream of successive areas, neurons gradually increase their receptive field sizes, become more selective to spatiotemporal stimulus patterns in their receptive fields (RFs), and also develop selective invariances (Felleman and Van Essen, 1991; Riesenhuber and Poggio, 2001).

A generally supported Mountcastle’s (1978) conjecture is that such progressive elaboration of feature tuning properties is accomplished by recursive application of essentially the same computational operation, performed by series of cortical columns on their afferent inputs (Phillips and Singer, 1997). However, the nature of this hypothesized operation, as well as the nature of the extracted features are poorly understood. Neurons acquire their mature feature-tuning properties to a large degree by learning from experience, using lower-level features provided by their afferent inputs to build higher-level ones. Technically, a feature is a mathematical transfer function over a set of afferent inputs to a neuron (or to a node in an artificial neural network). Each neuron has to select (learn) some useful transfer function. However, this can be a challenging task. In high-level cortical areas that are closely engaged in shaping the behavior, the neurons’ tuning to stimulus features can, in principle, be guided directly by their more or less obvious behavioral utility. But for early sensory areas, the identity of low-level stimulus features that would be behaviorally useful – as building blocks enabling the construction of high-level behaviorally significant features – and thus worth extracting is far from clear. Such low-level features might be too far removed from actual behavior for a criterion of “behavioral usefulness” being of practical use in their selection, at least initially during early postnatal cortical development. Instead, selection of such features would have to rely on some local signs promising their eventual usefulness.

The prevailing consensus, which emerged in the 1990s, is that such local signs of tentative features’ potential usefulness can come from the spatial and/or temporal context in which these features occur (Barlow, 1992; Becker and Hinton, 1992; Becker, 1996; Stone, 1996; de Sa and Ballard, 1998; Phillips and Singer, 1997; Hawkins, 2004; Favorov and Ryder, 2004). According to this idea, local but ultimately behaviorally useful features should be the ones that can be predictably related to other such features, either preceding or following them in time or taking place side-by-side with them. Thus, neurons should choose features for their ability to predict and be predictable from other such features. Predictive relations exist among features extracted from non-overlapping sensory inputs because they do reflect order present in the environment.

Thus, contextually predictable features are signatures of causal factors operating in the individual’s environment, which might be relevant to the individual’s interactions with its environment and therefore worth tuning to (Phillips and Singer, 1997; Ryder and Favorov, 2001; Favorov and Ryder, 2004; Ryder, 2004).

While this proposal is straightforward at the conceptual level, its actual algorithmic and neural implementational details – which ultimately establish its biological feasibility – are lacking and need fleshing out. In Section 2 of this paper, we use the known details of cortical functional architecture as guiding constraints to formulate a biologically realistic, algorithmically explicit computational model for contextually guided feature tuning in cortical columns. This model allows us to devise a version of multi-view canonical correlation analysis to explicitly extract shared contextual information from neighboring cortical columns, estimate its dimensionality, and compute the principal axes (basis vectors) of the space of contextually predictable features of input patterns occurring in a cortical column’s RF. In Section 3, we apply this methodology to natural images to reveal the information space of contextually predictable features available to a column in the primary visual area, V1. We train our cortical model in Section 3 on visual inputs obtained from natural images and demonstrate that the model’s neurons are highly capable of tuning to contextually predictable nonlinear features despite using only Hebbian synaptic plasticity. The model neurons learn close to all theoretically available contextually predictable features, and these features are found to be similar to those of neurons in the cat V1.

The demonstrated feature tuning effectiveness and biological realism of the model suggest that it might capture the core operation performed by cortical columns in their feedforward extraction of perceptually and behaviorally significant information. In a related paper (Kursun et al., 2024), we demonstrate that convolutional neural networks (CNN) trained using contextual guidance can perform better than deep CNN, which are trained using error-backpropagation, on visual and hyperspectral imaging tasks, tactile texture discrimination, or text classification.

## 2 THEORETICAL MODEL SPECIFICATION

### 2.1 Contextual guidance in cortical layer 3

Cerebral cortex is a complex dynamical system dominated by feedback circuits, but we limit our exploration to the feed-forward component of this system, which endows neurons with their identity-defining so-called “classical” RFs and feature-tuning properties. We further confine our exploration to the central pathway in the feed-forward elaboration of cortical neurons’ properties, which proceeds through a repeating sequence of two cortical layers. Cortical layer 4 (L4) is the principal initial recipient of the feed-forward afferent input to a cortical area. L4 converts that input into a new form and sends it, in particular, to layer 3 (L3) of the same cortical area for its feature-extraction operation. The product of that L3 operation is then sent to L4 of the next cortical area, where the same two-stage feature-extracting operation is repeated, but on a higher level, building on the advances made by the preceding cortical area (Rockland and Pandya, 1979; Felleman and Van Essen, 1991; Callaway, 2004).

In addition to their afferent input from L4, L3 neurons receive extensive contextual input via long-range horizontal connections from surrounding columns up to several millimeters away from their resident column (Gilbert and Wiesel, 1983; DeFelipe *et al*., 1986; Lund *et al*., 1993; Burton and Fabri, 1995). These contextual inputs are expected to guide L3 neurons to the sources of mutual information in these two, afferent and contextual, input sets (Phillips and Singer, 1997). Two principally different kinds of such sources are possible. First, the two input sets might have partially overlapping RFs; in other words, they might share some neurons along their afferent pathways in common. Such an *internal* source of mutual information in the two sets is trivial and has to be avoided. Second, the afferent and contextual input sets might be impacted by the same environmental agent. Such an *external* source of mutual information has the potential of being behaviorally significant and therefore worthy of recognition. One way to ensure that mutual information in the two input sets comes from external sources is to use inputs with non-overlapping RFs. Indeed, long-range horizontal connections come from far enough to have non-overlapping RFs but are close enough to reflect the same distal variables in the engaged environment.

The value of the feature (*φ*) extracted by the *i*^*th*^ L3 cell is computed from the afferent inputs to its resident cortical column *m*:

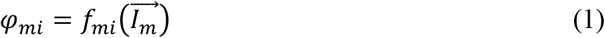

where *f*_*mi*_ is the chosen feature-specific transfer function, and 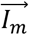 is a vector of activities of all the afferent axons innervating column *m*. According to the contextual guidance proposal, the function *f*_*mi*_ is chosen so as to maximize correlation of *φ*_*mi*_ with the “best” function *g*_*mi*_ over features extracted in other, surrounding columns and delivered via long-range horizontal connections:

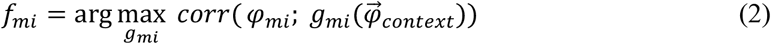

where *corr* is Pearson’s correlation coefficient, and 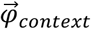 is a vector of all available L3 contextual features combined.

Since different neurons in a column should extract different features, we should elaborate the choice of the feature-extracting transfer function *f*_*mi*_ for a single neuron *i* in column *m*: the choice is to maximize correlation of *φ*_*mi*_ with the “best” function *g*_*mi*_ over features in other columns, subject to the constraint that correlation of this feature with features computed by other neurons in the *same* layer in the *same* column should not be excessive.

It should be noted that features extracted by cortical neurons – especially in high-level cortical areas – are highly nonlinear functions of peripheral patterns of receptor activations. No recursive application of linear transform functions would be able on its own to produce such features. Cortical neurons must be able to use nonlinear transfer functions. However, experience-driven learning of nonlinear transfer functions in neural networks can be highly problematic, unless a kernel-based strategy is used. Kernel methods, popular in machine learning, offer a highly effective strategy for dealing with nonlinear problems by transforming the input space into a new “feature” space where a nonlinear problem becomes linear and thus more tractable with efficient linear techniques (Schölkopf and Smola, 2002). According to Favorov and Kursun (2011), such a kernel-based function linearization strategy happens to be used in the neocortex in its principal input layer, layer 4. This insight suggests that cortical columns first perform a nonlinear function-linearization transform of their afferent inputs in L4 and then learn linear transform functions in L3.

### 2.2 Layer 4 pluripotent function linearization

An important feature of L4 functional architecture is the presence of untuned feed-forward inhibition, which reflects the overall strength of the stimulus activating a local L4 network but is insensitive – invariant – to spatial details of the stimulus patterns (Kyriazi et al., 1996; Bruno and Simons, 2002; Swadlow, 2003; Hirsch et al., 2003; Sun et al., 2006; Cruikshank et al., 2007). Favorov and Kursun (2011) showed that the presence of such untuned feed-forward inhibition converts a conventional neural network into a functional analog of Radial Basis Function (RBF) networks (Lowe 2003), which are well known for their universal function approximation and linearization capabilities (Park and Sandberg, 1991; Kurkova, 2003). Input transforms performed by such networks automatically linearize a broad repertoire of nonlinear functions over the afferent inputs. This capacity for pluripotent function linearization suggests that L4 can contribute importantly to cortical feature extraction by performing such a transform of afferent inputs to a cortical column that makes possible for neurons in the other layers of the column, including L3, to extract nonlinear features of afferent inputs using mostly linear operations.

A biologically realistic and highly effective pluripotent function linearizer has the following ingredients (Equation 3): (1) activity of each excitatory L4 cell is computed, in part, as a weighted sum of its afferent inputs, which are Hebbian; (2) lateral interconnections among L4 cells are used to diversify the afferent connectional patterns among L4 cells in a cortical column and give them a rich variety of RF properties; and (3) feed-forward inhibition makes L4 cells behave similarly to RBF units and is principally responsible for function linearization capabilities. Following Favorov and Kursun (2011), we describe L4 operation as:

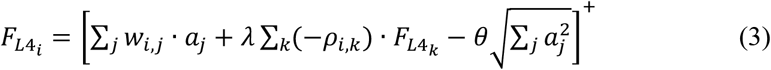

where 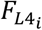 is the activity of L4 neuron *i*; *a*_*j*_ is the activity of afferent input neuron *j*; *w*_*i,j*_ is the weight, or efficacy, of the excitatory synaptic connection from afferent neuron *j* to L4 neuron *i*; *λ* is a lateral connection scaling constant; 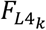 is the activity of a neighboring L4 neuron *k*; *ρ*_*i,k*_ is the correlation coefficient between activities of L4 neurons *i* and *k*; *θ* is a feed-forward inhibition scaling constant; and [·]^+^ indicates that if the quantity in the brackets is negative, the value is to be taken as zero.

In the presence of both feed-forward inhibition and plastic lateral connections, which are unique to L4 in that higher correlation in firings of the pre- and post-synaptic cells leads to decrease – rather than increase – of synaptic strength (Egger et al., 1999; Sáez and Friedlander, 2009), a modelled network of L4 neurons, trained on visual inputs, develops biologically accurate diversity of multi-subfield RFs and acquires orientation tuning matching in sharpness that of real L4 neurons, as well as a host of other real L4 functional properties (Favorov and Kursun, 2011).

In the above L4 model, neurons accomplish their pluripotent function linearization by acting in local groups. To explain, consider a local group of L4 neurons that are innervated by a common set of afferent neurons. Together, such a set of *N* convergent afferent neurons can be viewed as defining an abstract *N*-dimensional afferent input state space, each dimension corresponding to one of the constituent afferent neurons. A targeted L4 neuron *i* takes a particular direction in this afferent space (defined by the vector of its afferent connection weights 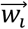 in Equation 3) as its conic RBF center. Neighboring L4 neurons innervated by the same set of afferent neurons chose different RBF centers (i.e., different 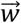) in their common afferent space, influenced in their choices by lateral interactions with each other. Together, a local group of L4 neurons will spread their RBF centers evenly throughout their common afferent space so as to map it most efficiently, with more active regions of that space mapped at higher resolution (Deco and Obradovic, 1995). A wide range of nonlinear functions defined over this space – including transfer functions that can extract contextually predictable features – can be then approximated by weighted sums of the activities of the mapping L4 neurons.

What is the size of such RBF-network-like local groups of L4 neurons that together can perform pluripotent function linearizations? Although it is limited, anatomical evidence based on typical sizes of afferent axon arborizations in L4 and the lateral spread of dendrites and axon collaterals of L4 cells within the confines of L4 suggests that such groups should be larger than a minicolumn (i.e., > 50μm in diameter) but smaller than a macrocolumn (i.e., < 500μm). We will refer to such intermediate-size function-linearizing columns in this paper as *mesocolumns*.

Structurally, minicolumns are the radially oriented cords of neuronal cell bodies evident in Nissl-stained sections of the cerebral cortex (Mountcastle, 1978; Buxhoeveden and Casanova, 2002). They are the narrowest (~50μm diameter) columnar aggregates of neurons in the neocortex (Favorov and Diamond, 1990; Tommerdahl et al., 1993) and thus can be viewed as the smallest building blocks of cortical columnar organization (Mountcastle, 1978). Published estimates of L4 cell densities in visual and somatosensory cortical areas (Beaulieu and Colonnier 1983; Budd 2000; Markram et al., 2015; Meyer et al., 2010) suggest that a single minicolumn has between 30-60 excitatory L4 neurons. Such a number is clearly not enough for a functionally useful RBF mapping of a minicolumn’s afferent space.

Minicolumns are packed together in the cortex in an essentially hexagonal pattern. From a geometric perspective, the next larger-size columnar entity to consider is a group of 7 minicolumns, one surrounded by 6 others. Such columns will be 3 minicolumns wide and thus ~150μm in diameter. They will have between 200-400 excitatory L4 neurons. In Section 3 we will show that such numbers of L4 neurons are sufficient for the purposes of contextually predictable feature extraction in L3. We propose that, based on the available evidence, such groupings of 7 minicolumns are the most plausible candidates for the role of function-linearizing mesocolumns (Figure 1). A group of 7 such mesocolumns, in turn, make the next larger-size columnar entity ~450μm in diameter, corresponding in size to the well-known macrocolumns (Mountcastle, 1997).

**Figure 1.**
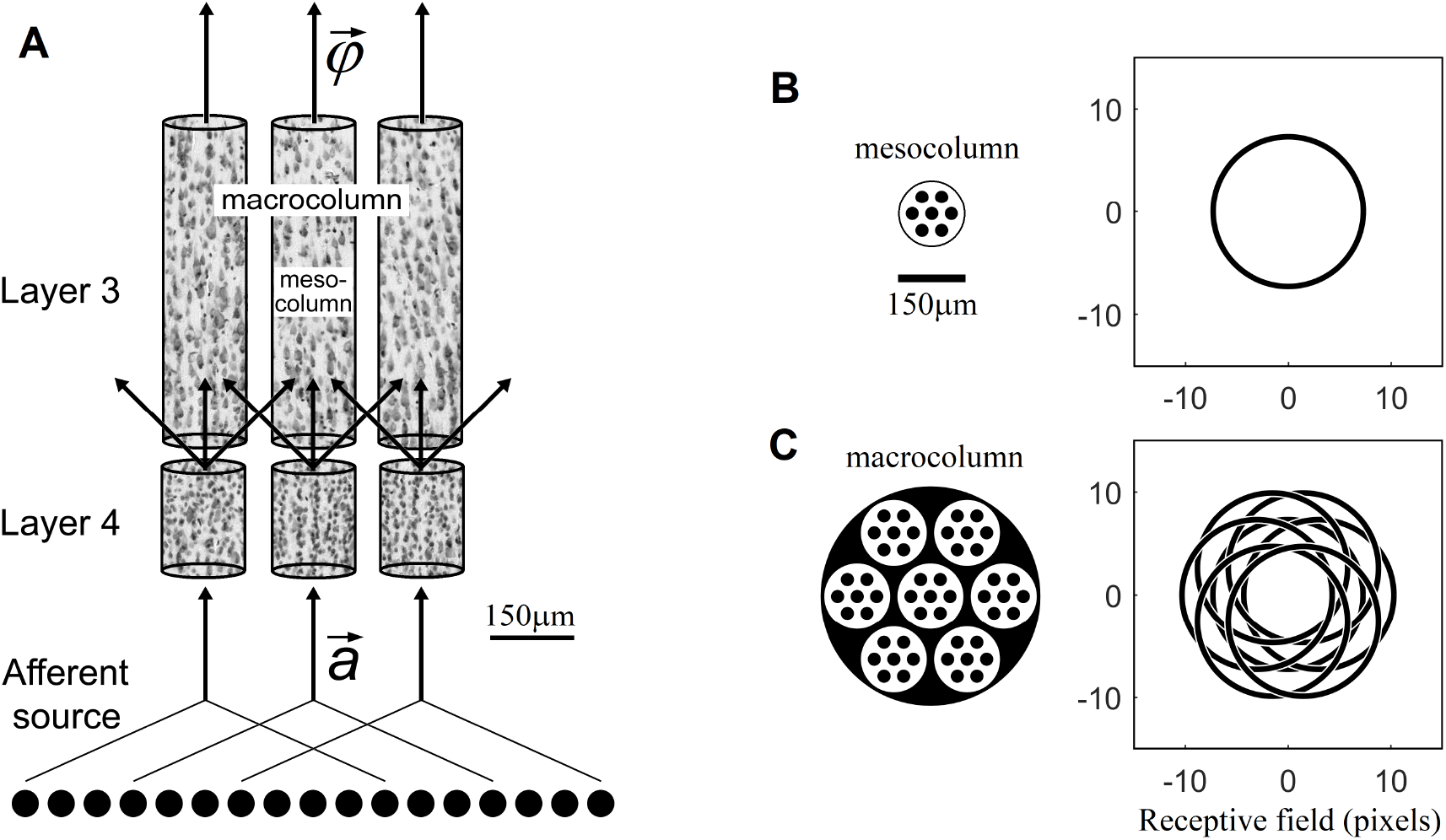
Mesocolumn-based feature extraction in the neocortex. **(A)** Connectional diagram. **(B)** Minicolumnar composition and RF of a mesocolumn. **(C)** Mesocolumnar composition and RF of a macrocolumn. The smallest structural units of neocortical columnar organization are *minicolumns*, comprising neurons whose bodies line up in ~50μm diameter radially oriented stacks separated by radially oriented bundles of axons and apical dendrites. Neurons residing in neighboring minicolumns are not functionally isolated but make up larger-size functional aggregates. This paper explores the feature extracting capabilities of aggregates that span local groups of 7 minicolumns, referred to as *mesocolumns*. The L4 and L3 compartments of three such mesocolumns, taken from monkey somatosensory cortex, are shown in panel **A**, each revealing multiple vertical stacks of Nissl-stained neurons. Neighboring mesocolumns receive their input 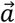 from partially overlapping sets of afferent neurons and thus from partially overlapping RFs. In panel **B**, a mesocolumn is shown schematically as a hexagonally packed group of 7 minicolumns (black filled circles). It is estimated to contain 200-400 excitatory cells in each of its L4 and L3 compartments. In the primary visual cortex, a mesocolumn’s RF in natural images would correspond to an approximately 16 pixel diameter circle (Favorov and Kursun, 2011). L4 cells in a mesocolumn act together as a group in performing pluripotent function linearizing transform of their RF input patterns. Neurons in the L3 compartment of a mesocolumn receive their afferent input from L4 cells residing not only in their own but also neighboring mesocolumns (panel **A**). Such a larger group of 7 mesocolumns, feeding central mesocolumn’s L3 neurons, is shown in panel **C** schematically as an ~450μm diameter *macrocolumn*. Since RFs of L4 compartments of these 7 mesocolumns are partially shifted (as shown), the overall RF of a mesocolumn’s L3 compartment is expanded to an approximately 21 pixel diameter circle. L3 neurons in a mesocolumn respond to diverse features of input patterns appearing in their mesocolumn’s RF, together converting the mesocolumn’s afferent input vector 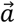 to the mesocolumn’s output feature vector 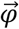.

### 2.3 Layer 3 extraction of contextually predictable features

The lateral spread of axonal projections of L4 cells in L3 and the lateral spread of basal dendrites of pyramidal cells in L3 indicate that L4 neurons send their output to L3 neurons not only in their own mesocolumn but also in the 6 surrounding mesocolumns (Lubke et al., 2003; da Costa and Martin, 2010). Consequently, the “classical” RFs and feature-tuning properties of L3 neurons in a given mesocolumn are, essentially, the product of weighted summation of output activities of L4 neurons of the same and 6 surrounding mesocolumns (Figure 1):

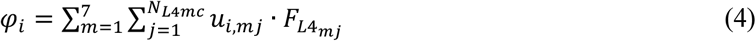

where *φ*_*i*_ is the feature-expressing activity of L3 neuron *i* in the central mesocolumn; *N*_*L4mc*_ is the number of L4 cells in a mesocolumn; 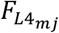 is the activity of an L4 neuron *j* in mesocolumn *m*; and *u*_*i,mj*_ is the strength of their connection.

In L3, the afferent inputs from L4 target basal dendrites of pyramidal cells, whereas the contextual inputs from surrounding cortical territories target their apical dendrites (Gilbert and Wiesel, 1983; Kisvarday et al., 1986; Lubke et al., 2003; Petreanu et al., 2009). Synaptic inputs to basal dendrites are integrated in the soma, leading to spike generation in the initial axon segment. But the apical dendrite, including its terminal tuft extension in layer 1, has its own site of synaptic input integration and is able to generate its own spikes (Bernander et al., 1994; Cauller and Connors, 1994; Schiller et al., 1997; Stuart and Spruston, 1998; Larkum et al., 1999, 2007). Output activity of the apical dendrite in the *i*^*th*^ L3 cell is, essentially, the product of weighted summation of output activities of L3 neurons of surrounding columns:

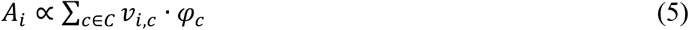

where *C* is the set of all the L3 neurons in surrounding columns that contribute contextual input to cell *i*; *v*_*i,c*_ is the strength of connection to cell *i* from contextual cell *c*; and *φ*_*c*_ is basal dendrite output of cell *c* (Equation 4). With such separate contextual input integration, the apical dendrite can guide basal dendrites in their selection of afferent connectional patterns (and vice versa) so that they will maximize covariance of the cell’s apical and basal outputs *A*_*i*_ and *φ*_*i*_, as was proposed and successfully demonstrated in a basic model by Kording and Konig (2000).

The number of excitatory cells in L3 is approximately the same as in L4 (Markram et al., 2015; Meyer et al., 2010), suggesting that the L3 compartment of a mesocolumn contains between 200-400 pyramidal neurons. Under mutual competitive pressure to diversify their RF tuning properties, similar to plastic local lateral connections among neighboring L4 cells driving them to select different features, these 200-400 neurons in a mesocolumn will compete in their search for contextually predictable features. Together, they will find and tune to all the different contextually predictable features present in their shared afferent input from L4 (Equation 4).

In geometric terms, together the 1400-2800 L4 cells in 7 mesocolumns that provide afferent input to the L3 compartment of a central mesocolumn create that mesocolumn’s high-dimensional *state space*. Since we are concerned with L3 features that are computed linearly in that state space (Equation 4), such features correspond to particular directions in the mesocolumn’s L3 state space: for a given L3 neuron *i*, its afferent connectional vector 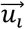 determines that neuron’s preferred direction in its mesocolumn’s L3 state space and thus its preferred feature.

Any arbitrary direction in the L3 state space will express some feature of input patterns taking place in the mesocolumn’s RF. However, most of such arbitrarily chosen features will not be perceptually significant. Contextually predictable features occupy a lower-dimensional subspace of the L3 state space. The size and contents of this contextually predictable subspace have never been revealed before, even for the primary sensory cortical areas. In the next section (Section 2.4), we derive a computational algorithm for estimating the principal axes (basis vectors) of this subspace, and in Section 3 we apply this algorithm to natural images in an attempt to reveal the contextually predictable subspace of a mesocolumn in the primary visual area, V1. Also in Section 3 we investigate what features (i.e., directions in the mesocolumn’s L3 state space) individual L3 cells will choose if they are modeled as comprising two dendritic compartments – basal and apical, each receiving Hebbian connections from either L4 of its own macrocolumn or from L3 of surrounding macrocolumns, respectively – and are trained on natural images. We show that such modeled L3 cells do indeed select features in the mesocolumn’s contextually predictable subspace.

### 2.4 Extraction of the contextually predictable feature subspace of a cortical mesocolumn

We begin by formalizing terminology to be used in the rest of this paper:

- The *state space* of a mesocolumn is created by 1400-2800 L4 cells that provide afferent input to its L3 compartment (Equation 4). Stimulus patterns activating that mesocolumn’s RF are represented as points in the state space and can be characterized in an infinite number of ways by projecting these points onto any particular vector in the state space. Any such projection vector can be considered a “feature,” and the entire state space of a mesocolumn is the space of possible features, or the feature space. Thus, we can refer to a mesocolumn’s state space as its *feature space* to emphasize its feature content.
- We search for contextually predictable features because such features reflect orderly aspects of the environment. To emphasize their orderly nature, we will follow Hotelling (1936) and call them *canonical features* (from the Greek word *kanonikotita*, κατ η τ ακóτητα, which means “regularity,” “predictable recurrence”). The contextually predictable subspace of a mesocolumn’s state space, comprising all the canonical features, then is the *canonical feature subspace*.
- Our explicit task is to extract the canonical feature subspace from the mesocolumn’s state space by finding all of its principal axes (basis vectors). This set of orthogonal vectors in the state space, enclosing the canonical feature subspace, will be called *canonical variates* (Hotelling, 1936).

We formulate our approach based on the following considerations. We define the 1^st^ axis of the canonical feature subspace (i.e., the 1^st^ canonical variate) to be the basis vector with the maximal correlation with the contextual input, the second axis (i.e., the 2^nd^ canonical variate) to be the basis vector with the second largest correlation with the contextual input, and so on until the last axis. In the cortex, different mesocolumns develop their own sets of afferent, lateral, and contextual connections based on their particular histories of sensory experiences. However, since neighboring mesocolumns will end up being exposed to and being shaped by the same regularities in their sensory experiences, any emergent differences among them will not be functionally significant. Thus, in deriving our algorithm, we can make an assumption that all the neighboring mesocolumns involved in contextual guidance will have the same matrices of L3 afferent [*u*_*i,mj*_] and contextual [*v*_*i,c*_] connections (Equations 4 and 5) and also identical sets of canonical variates.

To quantify the correlation of a canonical variate with the contextual input, we use the mean of pairwise correlations of that variate in the central mesocolumn and the same variate in each of the neighboring mesocolumns that contribute the contextual input. If we label the direction of a canonical variate in the L3 state space as 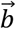 (*b* stands for “basis”), then we define the contextual correlation of this variate as:

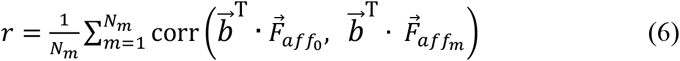

where *N*_*m*_ is the number of mesocolumns contributing contextual input. 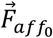 and 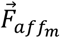 are the afferent inputs to the L3 compartment of the central (0^*th*^) mesocolumn and the *m*^*th*^ neighboring mesocolumn, respectively, from their flattened 7 × *N*_*L*4*mc*_ dimensional vectors of the outputs of L4 neurons of the same and 6 immediately surrounding mesocolumns:

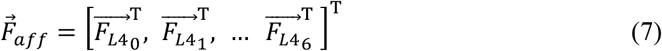

Our objective function for the *i*^*th*^ canonical variate is to find such a direction 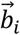 in the L3 state space that will maximize its contextual correlation *r*_*i*_ (subject to the constraint that 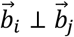 for all *j* < *i*). Our objective function is designed to maximize the canonical correlation of the *i*^*th*^ component, concurrently ensuring orthogonality with all previously computed components. This methodology, which constructs orthogonal vectors sequentially, beginning with the first, systematically generates a series of orthogonal vectors. Each vector maximizes the variance subject to the orthogonality constraints imposed by its predecessors.

Continuing with our assumption that different mesocolumns in a contextually related cortical territory have the same internal connectivities, we also assume that all *N*_*m*_ mesocolumns in our model have the same means and covariance matrices of the afferent inputs to their L3 compartments:

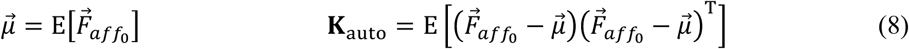

Then, we can write our objective function as:

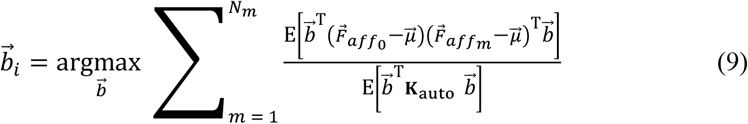

Thus, the objective function in Lagrangian formulation is given by:

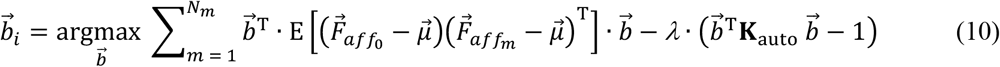

The optimization task specified in Equation 10 has a well-structured generalized eigenproblem and can be efficiently solved using established numerical algorithms (Hotelling, 1936; Hardoon, 2004; Kursun et al., 2011; Golub and Van Loan, 2013; Alpaydin, 2014), in which the eigenvector having the largest eigenvalue giving us the first canonical variate 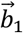, the eigenvector having the second largest eigenvalue giving us the second variate 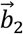, and so on:

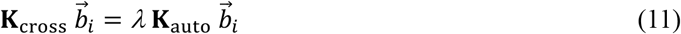

where K_cross_ is the cross-covariance matrix (Hotelling, 1936; Kursun et al., 2011):

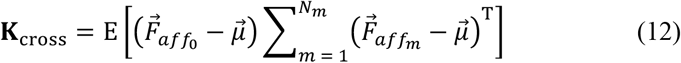

## 3 MODEL SIMULATIONS

For model simulations, we sought to emulate the primary visual cortical area (V1) and so applied our L4-L3 feature-extracting model to natural images, setting the afferent inputs of the modeled mesocolumns to approximate thalamic inputs to V1 from the lateral geniculate nucleus. The aim of simulations was to develop contextually predictable features that can be expected to be found in a representative V1 mesocolumn. Visual input patterns and the L4 compartments of modeled mesocolumns were reproduced, with a few minor differences, from Favorov and Kursun (2011) and that paper should be consulted for their complete descriptions.

## 3.1 METHODS

### 3.1.1 Visual input patterns to the L4 compartment of a mesocolumn

Biologically realistic visual afferent inputs, delivered to L4 via the lateral geniculate nucleus (LGN), were simulated based on the retinal/LGN model of Somers et al. (1995). RFs of LGN neurons were modeled as a difference of the “central” and the “surround” two-dimensional Gaussians, with a common space constant *σ* for both dimensions:

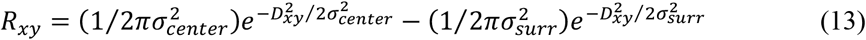

where *σ*_*center*_ = 0.8833 and *σ*_*surr*_ = 2.6499 (Figure 2A). *D*_*xy*_ is the Euclidean distance between a pixel at the (*x, y*) location in the image and the (*x*_0_, *y*_0_) image location of the RF center. If *D*_*xy*_ > 8, *R*_*xy*_ = 0 (i.e., the RF diameter is restricted to 16 pixels). Thus, the activity of an ON-center LGN neuron with the RF center at the (*x*_0_, *y*_0_) location in the image was computed as:

**Figure 2.**
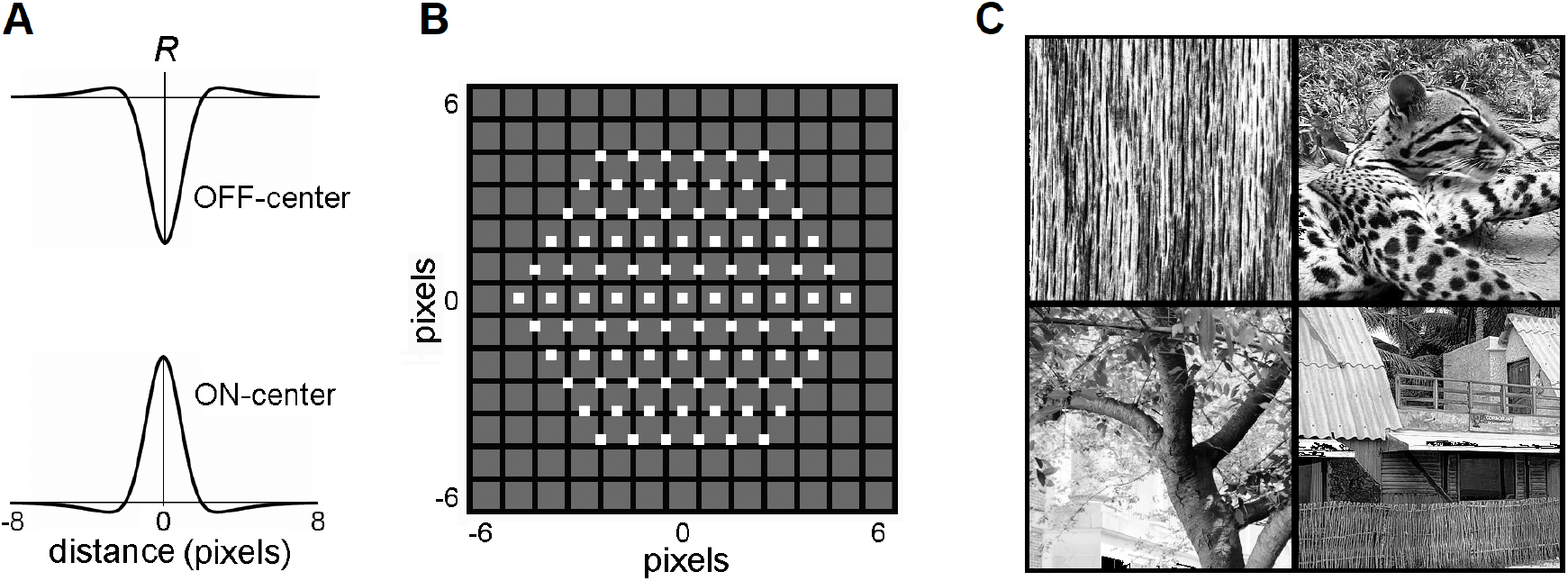
LGN layer model. **(A)** RF profiles of ON-center and OFF-center model LGN cells. **(B)** The map of the RF centers (little white boxes) of the 91 ON-center cells of the LGN layer innervating a single mesocolumn. Note that RF centers are arranged in a hexagonal pattern. RF centers are shown superimposed on a 13**×**13-pixel field, in which each pixel is shown as a black-edged gray square. RF centers of the 91 OFF-center LGN cells match the RF centers of the ON-center cells. **(C)** Four exemplary 320**×**320-pixel natural images that were used to activate the LGN layer. Reproduced with permission from Favorov and Kursun (2011).

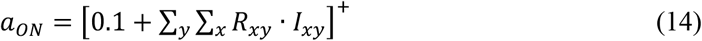

where *I*_*xy*_ is the grayscale intensity of the pixel at (x, y) location in the image (0 ≤ *I*_*xy*_ ≤ 1). The activity of an OFF-center LGN neuron was computed as:

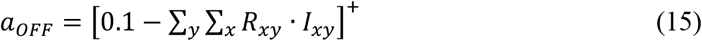

Each mesocolumn in the model was set to receive its afferent input from 91 LGN neurons with retinotopically arranged ON-center RFs and 91 neurons with retinotopically arranged OFF-center RFs (Figure 2B). RF centers of ON-center LGN neurons were arranged in a hexagonal pattern, spaced one pixel apart, and RF centers of OFF-center LGN neurons coincided with the RF centers of the ON-center LGN neurons. Together, these 182 LGN neurons created a hexagonally shaped viewing window onto visual images (Figure 2B).

In this study, the visual inputs to the LGN layer were drawn from a set of 100 grayscale photographs (320×320 pixels) selected from the IAPR TC-12 benchmark dataset (Grubinger et al., 2006), containing texture-rich natural images of surfaces, grass, bushes, landscapes, human and animal figures, and Brodatz (1966) dataset of textures (Figure 2C). Since this is a relatively small set of images, selected for their detail-rich spatial contextual information, an additional much larger and more diverse image dataset was also used to confirm the model findings made on the IAPR TC-12 dataset. This was a widely used *Common Objects in Context* (COCO) dataset of images of complex everyday scenes containing common objects in their natural context (Lin et al., 2015). In particular, we used 5000 images of the 2017 validation set (http://images.cocodataset.org/zips/val2017.zip HYPERLINK “http://images.cocodataset.org/zips/val2017.zip)”). The photographs were not preprocessed, except for contrast enhancement using histogram equalization. To generate a particular visual input pattern, the LGN viewing window was placed over a particular location in one of the photographs. The intensities of the pixels within the viewing window were then convolved with the RF profiles of the LGN neurons (Equations 13-15). All computational procedures were implemented using MATLAB (MathWorks, 2023).

### 3.1.2 Output of the L4 compartment of a mesocolumn

The L4 compartment of each mesocolumn was modeled as a group of *N*_*L4mc*_ neurons of the type described above by equation 3. The temporal behavior of each neuron, modeled as a leaky integrator, is described by the following differential equation:

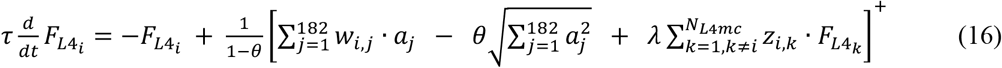

where τ is a time constant; 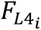 and 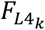 are the output activities of L4 neurons *i* and *k* in the computed mesocolumn, respectively; *a*_*j*_ is the activity of the *j*^th^ among the 182 LGN neurons innervating the computed mesocolumn; *w*_*i,j*_ is the strength of the afferent connection from LGN neuron *j* to L4 neuron *i*; *z*_*i,k*_ is the strength of the connection to L4 neuron *i* from L4 neuron *k* residing in the same mesocolumn; *θ* and *λ* are feed-forward and lateral connection scaling constants, respectively. This differential equation was solved numerically using Euler updates with a step size Δ*t* = 1ms. Explicitly, the Euler update for an equation *τ*(*d*/*dt*)*x* = −*x* + *g*(*x*) is *x*(*t* + Δt) ≈ (1 − Δ*t*/*τ*) · *x*(*t*) + (Δ*t*/*τ*) · *g*(*x*). Time constant τ was set to 4ms, *θ* = 0.65 and *λ* = 3. The response of the L4 network to a given afferent input pattern was computed in 20 time steps.

### 3.1.3 Hebbian development of afferent and lateral connections in the L4 compartment of a mesocolumn

The complete set of instructions and explanations offered in Favorov and Kursun (2011) should be followed in growing L4 connections. Partially repeated here, those connections were driven to their final state by modifying them iteratively over 20 update steps. At each step, the L4 compartment of a mesocolumn was stimulated with 1000 visual input patterns, which were produced by placing the LGN viewing window in random locations in any of the 100 database images. Output activities of the 182 LGN cells and *N*_*L4mc*_ L4 cells in response to these 1000 visual patterns were used to compute correlation coefficients between all pairs of LGN-L4 and L4-L4 neurons, and those correlations were used to update the afferent and lateral connections.

At update step *s*, the strength of the afferent connection from LGN cell *k* to L4 cell *i* was updated based positively on correlation *ρ*_*ik*_(*s*) of their outputs during step *s* as:

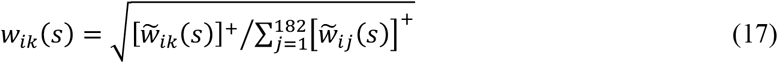

Where

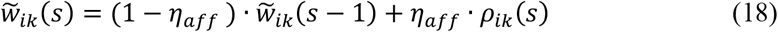

The weight of the lateral connection between L4 cells *i* and *k* was updated based negatively (according to Egger et al., 1999; Sáez and Friedlander, 2009) on correlation *ρ*_*ik*_(*s*) of their outputs during step *s* as:

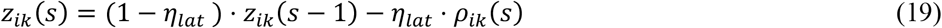

Adjustment rate constants *η*_*aff*_= 0.01 and *η*_*lat*_= 0.1 produced the fastest convergence of connection strengths to stable values.

### 3.1.4 Afferent and contextual inputs to the L3 compartment of a mesocolumn

The L3 compartment of a mesocolumn in the model was set to receive afferent input from its own L4 compartment as well as from L4 compartments of six immediately adjacent mesocolumns (Equation 4). The LGN viewing windows of these six surrounding mesocolumns are shifted by 3 pixels relative to the LGN window of the central mesocolumn in six evenly spaced directions (Figure 1c). Consequently, the viewing window of the L3 compartment of a mesocolumn (which we consider to be its classic RF) is increased to 21 pixels.

In the real cortex, L4 compartments of different mesocolumns develop their own sets of LGN and lateral connections based on their visual experiences. However, since visual experiences of 7 neighboring mesocolumns are essentially the same, any emergent connectional differences among them will not be functionally significant. This allows us to greatly reduce the computational effort in developing the model’s L4 connectivity by developing LGN and lateral connections of just one mesocolumn and then use these patterns of connections (i.e., the [*w*_*ij*_] and [*z*_*ik*_] matrices in Equation 16) in all the mesocolumns making up the model.

Our definition of the mesocolumn in Section 2.2 as a local group of 7 minicolumns, L4 cells of which together perform pluripotent function linearization transform of their shared afferent input, leads us to treat mesocolumns in this modeling effort as discrete entities packed in the cortex as a honeycomb-like mosaic. We also treat macrocolumns as discrete entities comprising 7 mesocolumns (Figure 3). However, this might be oversimplification. While discrete macrocolumns do exist – at least in the somatosensory cortex (Favorov and Diamond, 1990; Favorov et al., 2012) – experimental evidence of discrete mesocolumnar structures in L4 is lacking. It is possible that discrete mesocolumns, while appealing in their conceptual simplicity, are not necessary, and L4 function linearization transform can be successfully performed by a field of partially overlapping mesocolumns (making a mesocolumn a functional, rather than structural, entity). We will leave exploration of this possibility for future studies.

**Figure 3.**
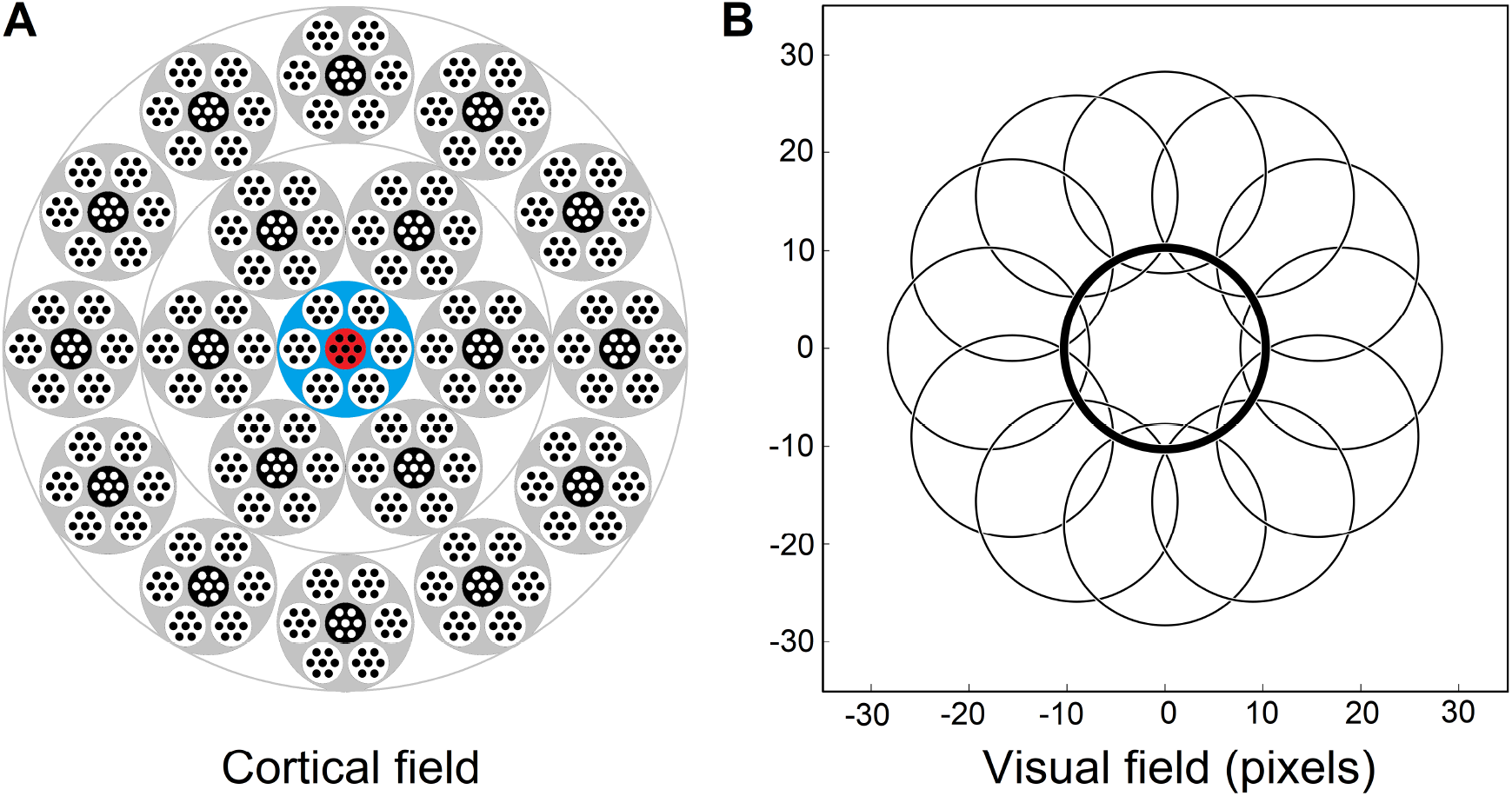
Afferent and contextual inputs to the model mesocolumn’s L3 compartment. **(A)** The central macrocolumn (blue shaded) surrounded by two concentric rings of 6 and 12 macrocolumns (gray shaded) carrying contextual information. In the cortex, these rings would be ~0.5 and ~1.0mm away from the central macrocolumn. In the model simulations, image-response activities were computed for L4 cells in all 7 mesocolumns in each of the 19 macrocolumns but, to reduce the amount of computation, responses of L3 canonical variates were computed only for the central mesocolumn in each macrocolumn. They were used as the contextual input to the L3 cells in the central macrocolumn’s central mesocolumn (red shaded). **(B)** RF outlines of the central and 12 outermost surrounding macrocolumns, showing very limited overlap. RFs of the inner and outer rings are shifted by 9 and 18 pixels, respectively, relative to the RF of the central macrocolumn.

Thus, for model simulations, the afferent input to the L3 compartment of the central mesocolumn is a flattened 7 × *N*_*L*4*mc*_ dimensional vector of the outputs of L4 neurons of the same and 6 surrounding mesocolumns (Equation 7). The contextual input to the L3 compartment of the central mesocolumn in the model was set to come from L3 compartments of two concentric rings of distant mesocolumns: the inner ring of 6 mesocolumns and the outer ring of 12 mesocolumns (Figure 3A). RFs of the outer ring mesocolumns are shifted by 18 pixels relative to the RF of the central/recipient mesocolumn (Figure 3B). RFs of the inner ring mesocolumns are shifted by half of this distance; i.e., by 9 pixels.

### 3.1.5 Algorithmic extraction of canonical variates

Canonical variates are the principal axes of the canonical feature subspace. To find their directions 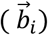 in the L3 state space (Equations 9-11), the modeled field of L3 compartments of 19 mesocolumns, each receiving afferent input from L4 compartments of its own and 6 surrounding mesocolumns (Figure 3), was stimulated with 5000 visual input patterns, which were obtained by placing the LGN viewing window in random locations in any of the 100 database images. For each visual pattern, the afferent input to the L3 compartment of each of the 19 mesocolumns in the field 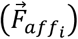 was written as a flattened vector of output activities of cells in L4 compartments of its own and its 6 surrounding mesocolumns (Equation 7). 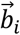 vectors of the first 20 canonical variates were extracted from the 5000 sets of afferent input vectors of the central and 18 surrounding mesocolumns 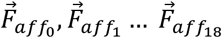 according to Equations 8, 11, 12. Using 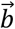 vectors, magnitudes of canonical variates of L3 afferent input patterns can be computed as:

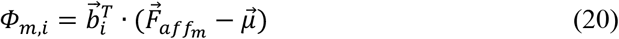

where *Φ*_*m,i*_ is the projection of the L3 afferent input vector of the *m*^*th*^ mesocolumn onto the *i*^*th*^ canonical variate.

### 3.1.6 Hebbian tuning of L3 cells to canonical features

The central thesis of this paper is that pyramidal neurons in L3 should tune to contextually predictable, *canonical*, features, and they accomplish it by adjusting the weights of L4 connections to their basal dendrites under guidance from their apical dendrites, which receive contextual inputs from the surrounding cortical territory (Section 2.3). To explore what features might be thus selected by L3 cells in a V1 mesocolumn, we gave the L3 compartment of the central mesocolumn the same number of cells as in its L4 compartment (i.e., *N*_*L3mc*_ = *N*_*L4mc*_) and trained their L4 input connections using an approach adapted from Kording and Konig (2000). For contextual guidance, we used canonical variates in the surrounding 18 mesocolumns (black shaded mesocolumns in Figure 3A).

Thus, the LGN viewing window was placed in 5000 random locations in the 100 database images, and for each image location we computed afferent input vectors to L3 compartments of the central and 18 surrounding mesocolumns 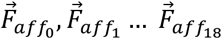 as well as the responses of canonical variates *Φ*_*m,i*_ to these input vectors (Equation 20). These responses were autoscaled to zero mean and unit variance.

The feature-expressing basal outputs of L3 cells in the central mesocolumn in response to images were computed according to Equation 4 while the contextual inputs to the apical dendrites of the same L3 cells were computed as:

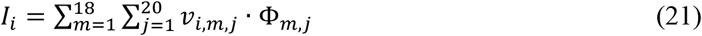

where *I*_*i*_ is the net contextual input to the apical dendrite of the *i*^*th*^ L3 cell, *Φ*_*m,j*_ is the response of the *j*^*th*^ canonical variate in the *m*^*th*^ surrounding mesocolumn, and *v*_*i,m,j*_ is the strength of their connection. Both basal outputs of L3 cells and net inputs to their apical dendrites were autoscaled to zero mean and unit variance.

The L3 network has to have a mechanism for diversifying feature tuning properties of cells residing in the same mesocolumn. The presence of such mechanism is indicated by the fact that in the real cortex, while neighboring neurons do share some of their RF and feature tuning properties in common, when all of these properties are considered *in toto*, neighboring neurons are very distinct and are highly decorrelated in their responses to the full repertoire of natural stimuli (Favorov and Kelly, 1996ab; Vinje and Gallant, 2000). The nature of this diversifying mechanism has not been established yet, but it must involve individual L3 cells in a mesocolumn influencing (likely “pushing” via lateral inhibition subserved by double-bouquet (DeFelipe et al., 2006) and/or Martinotti cells (Silberberg and Markram, 2007)) each other to select features different from their own. In the absence of the established mechanism, we chose to achieve its effect by using the same diversifying mechanism we (Favorov and Kursun, 2011) proposed to operate in L4.

Thus, to promote tuning of L3 cells in the mesocolumn to different canonical features, the contextual inputs to their apical dendrites were modified by the basal outputs of all the other L3 cells in the mesocolumn, as well as by the output of the mesocolumn’s L3 feed-forward inhibitory cell. That is, the output of the apical dendrite of the *i*^*th*^ L3 cell in the mesocolumn was computed as:

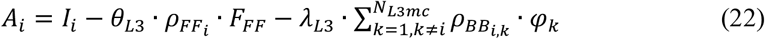

where *θ*_*L*3_ and *λ*_*L*3_ are feed-forward and lateral scaling constants; *φ*_*k*_ is the basal output of the *k*^*th*^ L3 cell and 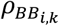 is the correlation between *φ*_*i*_ and *φ*_*k*_ over the training set of images; *F*_*FF*_ is the output of the L3 feed-forward cell and 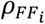 is the correlation between *φ*_*i*_ and *F*_*FF*_. *F*_*FF*_ was computed as the sum of outputs of all L4 cells in the central macrocolumn. It was autoscaled to zero mean and unit variance over the training set of images. The values of *θ*_*L*3_ and *λ*_*L*3_ scaling constants were tested systematically for their feature diversification effect on L3 cells by measuring cross-correlations between basal outputs of different L3 cells in the mesocolumn. Gradually increasing the values of these constants leads to gradual reduction of cross-correlations, starting from very high values to eventually very low, which indicate that different L3 cells tuned to different features. Based on this empirical search, the optimal settings of *θ*_*L*3_ = 0.01 and *λ*_*L*3_ = 0.03 were chosen, because under them L3 cells tune to the most diverse set of canonical features.

Hebbian connections of the basal and apical dendrites of L3 cells were developed gradually by modifying them iteratively over 1000 update steps. At each update step, the modeled field of 19 macrocolumns was stimulated with 5000 visual input patterns, which were obtained by placing the LGN viewing window in random locations in any of the 100 database images. Output activities of L4 and L3 cells and canonical variates in response to these 5000 visual patterns were used to compute correlation coefficients of L3 cells with L4 cells and with canonical variates, and those correlations were used to update the afferent and contextual L3 connections.

At update step *s*, the strength of the afferent connection from the *j*^*th*^ L4 cell to the basal dendrite of the *i*^*th*^ L3 cell was updated based on correlation of the L4 cell with the apical output, 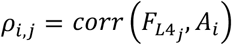, during 5000 step *s* trials as:

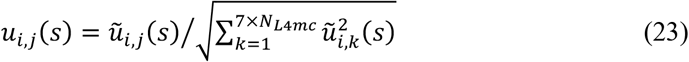

where

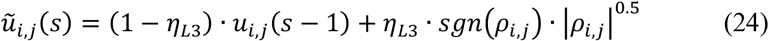

The strength of the contextual connection from the *j*^*th*^ canonical variate to the apical dendrite of the *i*^*th*^ L3 cell was updated based on correlation of the variate with the basal output, *ρ*_*i,j*_ = *corr*(*Φ*_*j*_, *φ*_*i*_), during 5000 step *s* trials as:

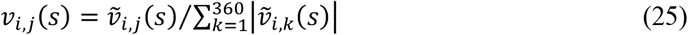

where

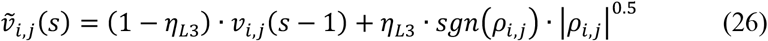

Adjustment rate constant *η*_*L3*_ = 0.01 produced the fastest convergence of connection strengths to stable values.

### 3.1.7 No-context L3 model

As an alternative to our contextually guided model of L3 feature tuning, we also tested a no-context model, in which the apical dendrite of each L3 cell was given exactly the same afferent input as its basal dendrite; i.e., instead of using Equation 21 to compute *I*_*i*_, we used *I*_*i*_ = *φ*_*i*_. That is, instead of receiving contextual input from surrounding columns, apical dendrites received their input from L4 cells of their own macrocolumn. Features developed by L3 cells in this essentially generic self-organizing neural network offer us a benchmark against which to judge the benefits of using contextual guidance in feature selection.

### 3.2 RESULTS

#### 3.2.1 Layer 3 canonical variates

According to our proposed division of tasks between L4 and L3 – with the L4 mesocolumnar network linearizing feature-extracting functions that will be computed by L3 cells – the first step in estimating L3 canonical variates is to develop RF and functional properties of cells in the L4 compartment of mesocolumns. This is done by repeatedly exposing L4 cells to images and adjusting the weights of their Hebbian input and intrinsic connections, gradually driving them into stable connectional patterns. The emergent functional RF properties of the model L4 cells, which come to closely resemble those of simple cells in cat V1, are comprehensively described in Favorov and Kursun (2011), and for brevity we omit their description here.

On their own, the trained L4 cells have very low pairwise contextual correlations with L4 cells in surrounding macrocolumns. This is shown in Figure 4A by plotting the distribution of maximal correlations of coincident activities of individual L4 cells in the central macrocolumn and L4 cells in the first and second rings of surrounding macrocolumns. However, our expectation is that optimally chosen weighted sums of multiple L4 cells will have much higher contextual correlations with surrounding macrocolumns. In Section 2.4 we introduced a particular algorithm for finding such optimal weighted sums, giving us the axes of the canonical feature subspace, i.e. canonical variates. We apply this algorithm to the outputs of L4 cells in the modeled field of 19 macrocolumns to obtain canonical variates. We test the strength of their contextual correlations by introducing a third ring of 12 mesocolumns, chosen to be at a such distance from the central mesocolumn that any mutual information they might have in their RFs will have to come from the environmental sources rather than from sharing any pixels in common. To compute their contextual correlations, we used responses *Φ* (Equation 20) of the canonical variates in the central and these 12 distant surrounding mesocolumns to 1000 image patches taken at random in the 100 dataset images. For each canonical variate, its contextual correlation is expressed by Pearson correlation coefficient computed between the 1000 responses of that variate in the central mesocolumn and the 1000 averages of responses of that variate in the 12 distant mesocolumns. The magnitudes of the computed contextual correlations are plotted in Figure 4B (white bars) for the first 20 canonical variates. The first 7 variates have particularly high correlations (*r*^2^ ≥ 0.1). Correlations of the 8^th^ to 15^th^ variates, although low, are nevertheless statistically significant (at α = 0.05 with Bonferroni correction), suggesting that even these canonical variates might reflect some causally significant factors in the environment.

**Figure 4.**
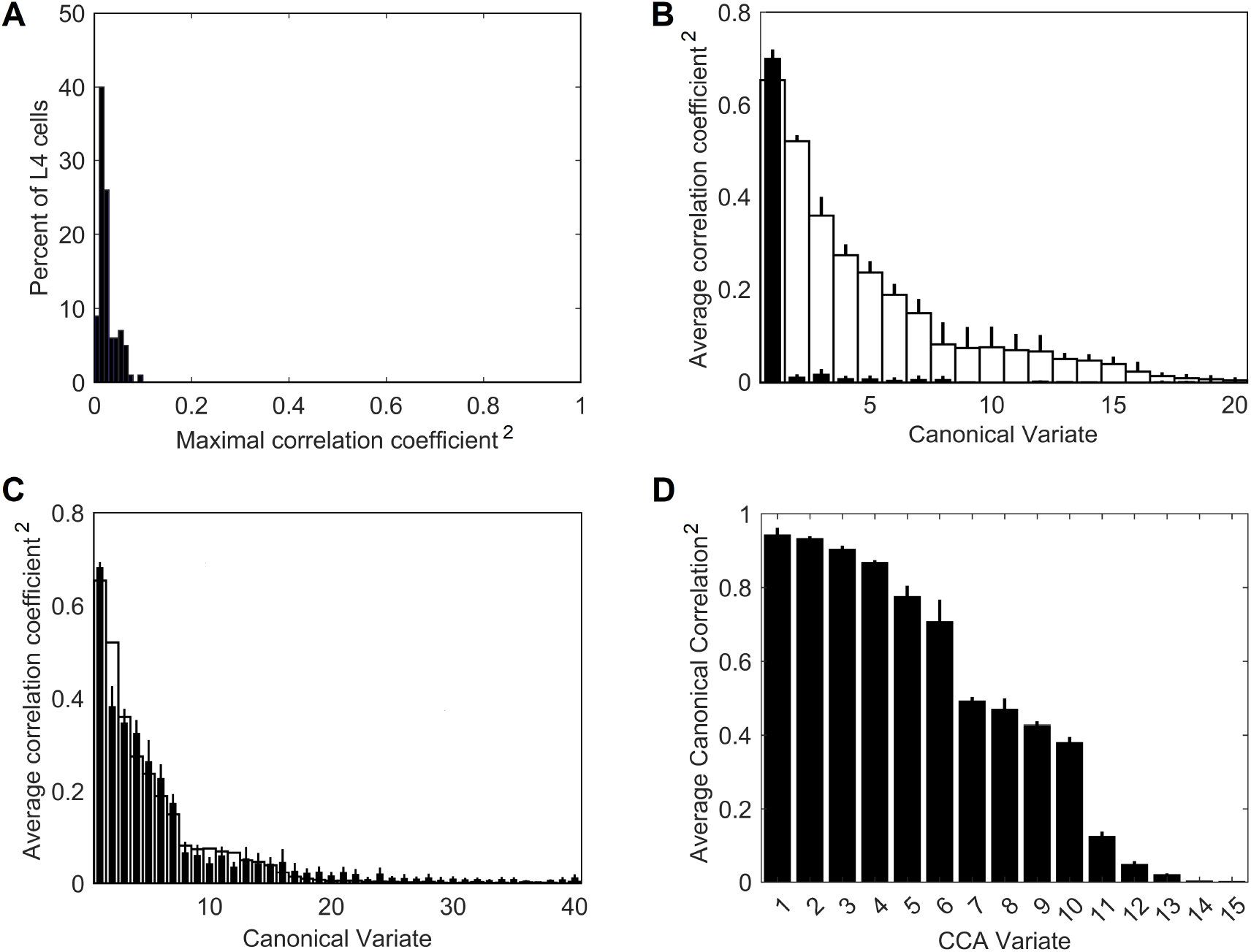
Contextual correlations between the central and surrounding macrocolumns. **(A)** Contextual correlations among cells in the input layer, L4. Plotted is the histogram of the highest correlation of stimulus-evoked responses that each L4 cell in the central macrocolumn had with L4 cells in the 2 rings of surrounding macrocolumns, revealing that at the level of L4, individual cells in neighboring macrocolumns were essentially uncorrelated. Each mesocolumn had 150 L4 cells. **(B)** Contextual correlations between the first 20 canonical variates of the central mesocolumn and the surrounding mesocolumns with nonoverlapping RFs. For each canonical variate, correlation was computed between its value in the central mesocolumn and the mean of its values in the surrounding mesocolumns. Training of L4 cells and extraction of canonical variates to find their contextual correlations was repeated 10 times, using different randomly chosen sets of training image patches taken from the 100 images of the IARP TR-12 and Brodatz datasets. Shown in the plot are squared correlation averages and their SEM (white bars), indicating that macrocolumnar RFs possess up to 15 canonical variates with contextually significant information. Also shown in the plot are squared correlation averages and their SEM of canonical variates extracted from LGN afferent inputs to macrocolumns (black bars), indicating that only the first LGN-based variate has significant contextual information. **(C)** Contextual correlations between the first 40 canonical variates extracted from the 5000 images of the COCO dataset (black bars). Shown in the plot are squared correlation averages and their SEM (n = 10). For comparison, also plotted superimposed are the first 40 canonical variates extracted from the 100 images of the IARP & Brodatz datasets (white bars), revealing close similarity between them. **(D)** Canonical Correlation Analysis (CCA) of overlap between canonical feature subspaces extracted by the first 15 canonical variates in the IARP & Brodatz vs. COCO datasets (details in the main text). Plotted are squared canonical correlations of the 15 CCA variates, averaged over doing CCA 10 times.

To demonstrate the necessity of the L4 function-linearization operation for maximizing contextual correlations, we also developed canonical variates directly from the LGN afferent inputs to the L4 compartments of the central and 6 surrounding mesocolumns (together constituting a macrocolumn) rather than using L4 outputs of these 7 mesocolumns. Unlike the L4-based variates, all but the first of the LGN-based variates showed no statistically significant contextual correlations (black bars in Figure 4B).

To test generalizable nature of the canonical variates extracted from the 100 IARP TC-12 images, canonical variates were also extracted from the 5000 COCO images. The magnitudes of their contextual correlations, shown for the first 40 variates, are plotted as black bars in Figure 4C, superimposed on the first 40 canonical variates extracted from IARP images (plotted as white bars). As Figure 4C shows, although they come from different sources, magnitudes of the two sets of canonical variates are very similar, with the first 15 variates having statistically significant contextual correlations. But how similar are the features extracted from the two image sources?

As basis vectors, the first 15 contextually predictable canonical variates enclose the canonical feature subspace of the mesocolumn’s entire feature space (as defined in Section 2.4). To determine how much the IARP and COCO canonical feature subspaces overlap, we performed Canonical Correlation Analysis (CCA; Hotelling, 1936), in which we treated the first 15 IARP and first 15 COCO canonical variates as 2 sets of input variables and used 5000 training image patches, taken at random from the COCO dataset, to compute their loadings. Next, we used these loadings to compute 15 canonical correlations of the two sets of variables over a different set of randomly picked 1000 COCO image patches. If the two feature subspaces, enclosed by the 15 IARP and 15 COCO canonical variates, match closely, the 15 canonical correlations would all be close to 1. On the other hand, if the two subspaces do not overlap at all, the 15 canonical correlations would all be close to zero. The actual computed correlations are plotted in Figure 4D, revealing that the first 6 CCA variates had very high correlations, whereas the last 5 CCA variates had very low correlations. Thus, we can conclude that canonical feature subspaces extracted from the IARP and COCO datasets mostly overlap, albeit not completely.

Going back to Figure 4B, as it shows, only the first canonical variate does not depend on L4 function-linearization preprocessing. The reason is that it reflects the overall magnitude of activity evoked in the macrocolumn’s L4 compartment (Figure 5A) and thus the overall stimulation intensity of the macrocolumn’s RF. Since the other canonical variates depend on L4 function-linearization preprocessing, they must be tuned to various structural features of the image patterns occurring in the mesocolumn’s RF. What these features are, either in our model canonical variates or in real L3 neurons, is not obvious but some insight is traditionally gained in V1 studies by characterizing responses of V1 neurons to moving grating images of various orientations and spatial frequencies. Figure 5B shows orientation and positional tuning of the statistically significant first 15 canonical variates, revealing that variates 8 through 11 are sensitive to both orientation and position while others are sensitive to grating orientation but not its position in the RF (translational invariance), thus falling into the categories of the *simple* and *complex* cells, respectively (Hubel and Wiesel, 1962). With real V1 neurons exhibiting diversity in the degrees of their orientation and grating phase tuning, the standard metric used to place any given V1 cell on the simple vs. complex cell spectrum is the F1/F0 ratio, which is the ratio of the 1^st^ and 0^th^ Fourier harmonics of a neuron’s activity during stimulation of its RF with an optimal sinewave moving grating (Skottun et al., 1991; Ringach et al., 2002). V1 cells with F1/F0 > 1 are classified as simple and cells with F1/F0 < 1 are classified as complex. Figure 5C shows F1/F0 scores of the statistically significant canonical variates 2-15, showing that 70% and 30% of variates fall into the complex cell and simple cell categories, respectively.

**Figure 5.**
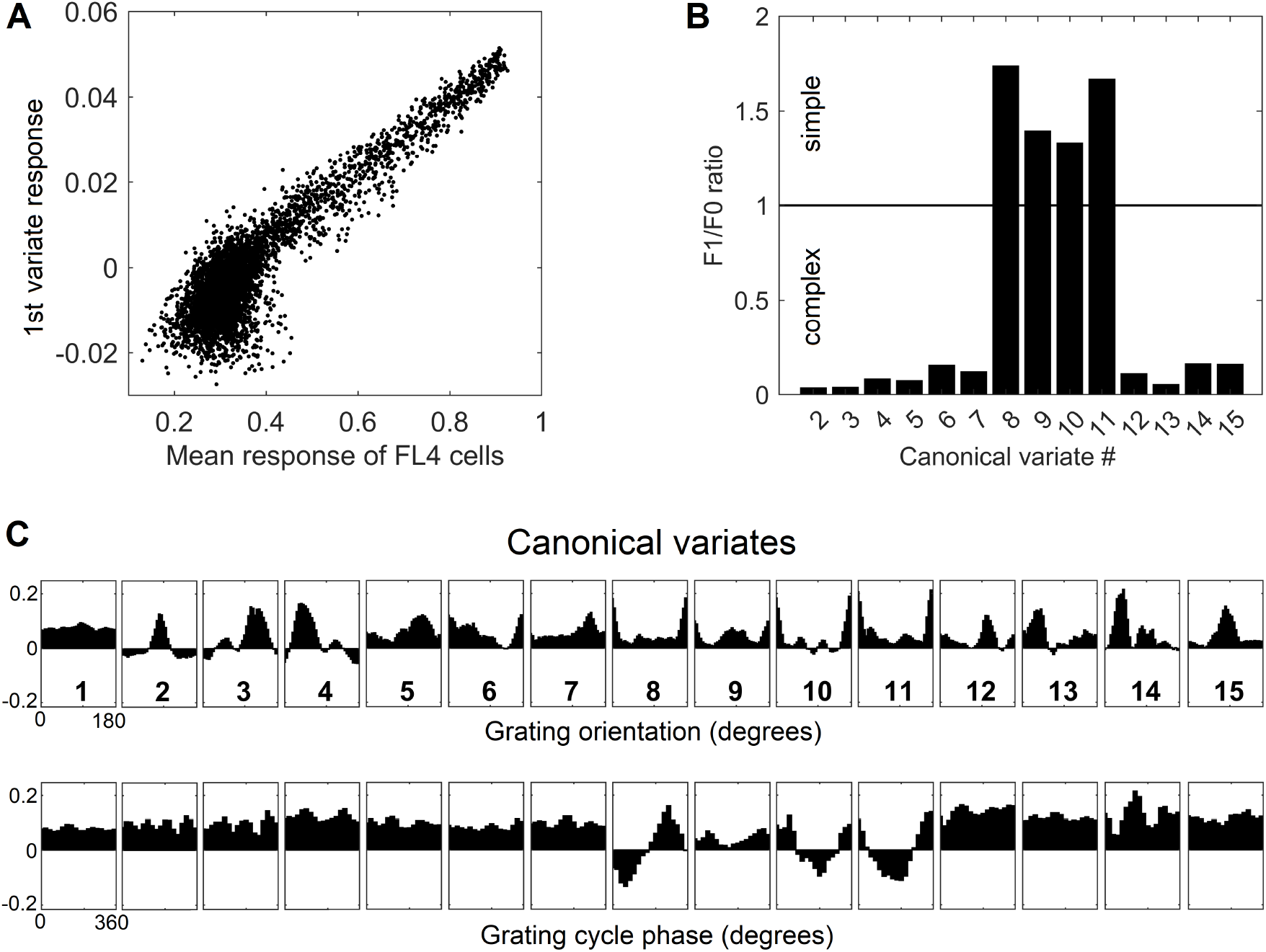
Feature tuning of canonical variates. **(A)** Tuning of the 1^st^ canonical variate to the overall intensity of RF stimulation. The variate’s response magnitude is plotted as a function of the average of the stimulus-evoked activities of all the L4 cells in the macrocolumn, showing linear dependency. **(B)** Tuning of the first 15 canonical variates to the orientation and spatial phase of sinewave grating images. **(C)** F1/F0 scores of canonical variates 2-15, showing clear separation of these variates into the simple and complex cell classes.

In principle, function-linearization capabilities of mesocolumns’ L4 compartment depend on the number of cells they employ (Favorov and Kursun, 2011): the larger the number of L4 cells in a mesocolumn, the broader the repertoire of nonlinear functions it can linearize. This is shown in Figure 6, in which the total contextual correlation of the first 20 canonical variates, computed as the sum of squared contextual correlations of individual variates (Watanabe, 1960), is plotted as a function of the number of cells in each mesocolumn’s L4 compartment. Significantly, there is little further gain in total correlation after the number of L4 cells in mesocolumns is increased beyond 150-200, which suggests that they linearize all the contextually predictable features available for extraction in the mesocolumn’s RF.

**Figure 6.**
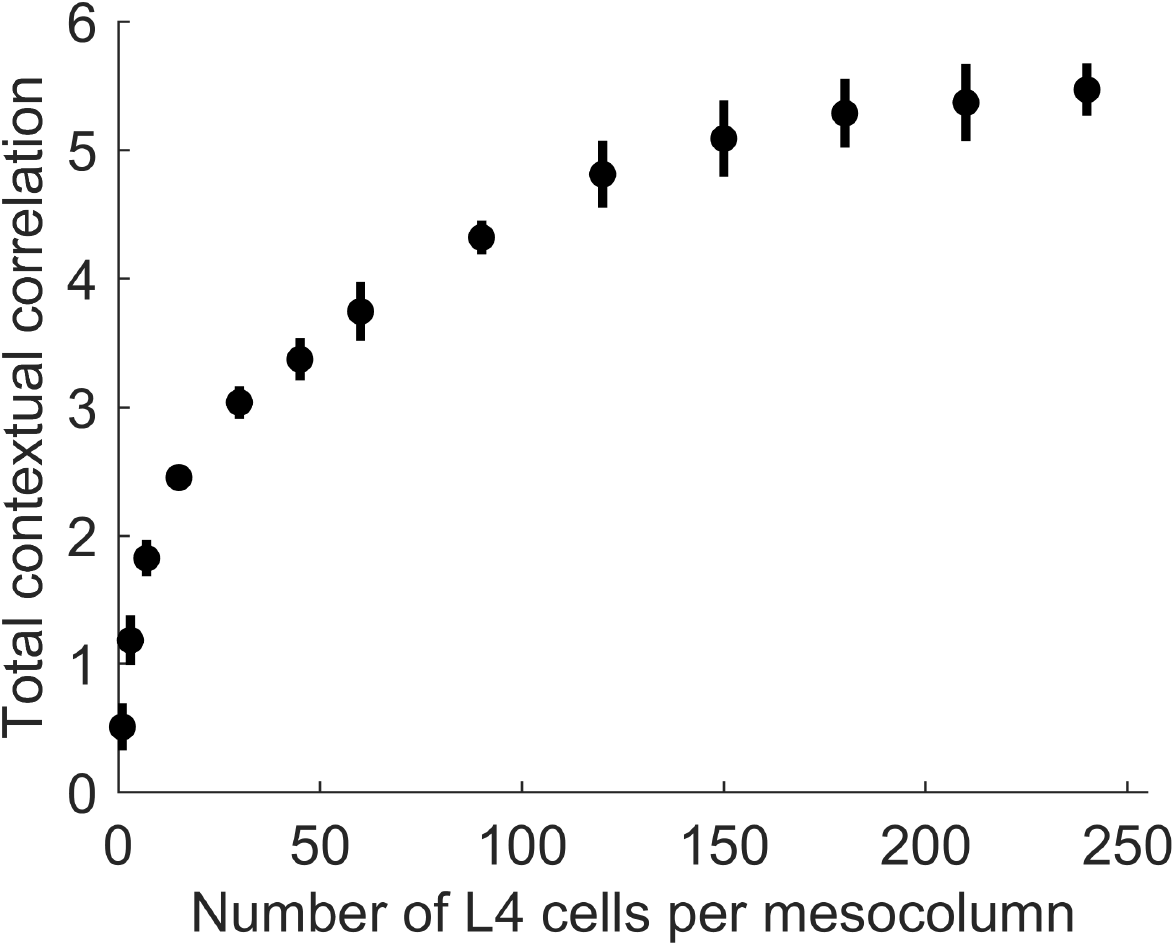
The dependence of the amount of contextually predictable information extracted by canonical variates on the number of L4 cells in the mesocolumn (*N*_*L4mc*_). The amount of contextual information extracted by the first 20 canonical variates was estimated by their total contextual correlation, which was computed as the sum of squared (with sign preserved) correlations of the 20 variates in the central mesocolumn and their averages among the 12 surrounding mesocolumns with abutting RFs. Total correlation was computed 10 times, each time using different randomly chosen sets of image patches to train a particular number of L4 cells per mesocolumn and to extract canonical variates. Plotted are the means and standard deviations of the total correlation estimated for the number of L4 cells per mesocolumn ranging from 1 to 240. The plot suggests that having around 150 L4 cells per mesocolumn might be enough to extract most of the contextually predictable information in mesocolumns’ RFs.

#### 3.2.2 Canonical features of L3 cells

L3 cells are expected to be driven by their apical dendrites to tune to contextually predictable – *canonical*, according to our terminology – features. Such features occupy a particular subspace in the mesocolumn’s state/feature space, and the extracted canonical variates give us the principal axes of this canonical feature subspace. In choosing their features, L3 cells should be attracted to the canonical variates according to their contextual predictability. We explored this feature-selecting mechanism in our extended field of 19 macrocolumns (Figure 3A) by providing the L3 compartment of the central mesocolumn (red-shaded in Figure 3A) with 150 cells, each modeled as a pair of basal and apical dendrites. In each L3 cell, its basal dendrite was given afferent input from 1050 L4 cells of its own and 6 immediately adjacent mesocolumns (150 L4 cells per mesocolumn), which together make up a macrocolumn (blue-shaded in Figure 3A). The vector of the weights of these L4 connections to the cell’s basal dendrite (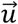 in Equation 4) determines that dendrite’s preferred direction in the mesocolumn’s L3 state space and thus that cell’s preferred feature. The apical dendrite of each L3 cell was given contextual input from the 18 more distant mesocolumns (black-shaded in Figure 3A) in the form of their 15 statistically significant canonical variates (for further details, see Methods section 3.1.6). The apical dendrite learns to produce output that best matches the output of the basal dendrite and vice versa.

Feature selectivities of L3 cells were developed by repeatedly exposing the model to randomly picked dataset images and adjusting the weights of the Hebbian basal and apical connections of L3 cells, gradually driving them into stable connectional patterns (Figure 7). After such training, the apical and basal dendrites of L3 cells developed prominent correlations in their responses to image patches (Figure 8A), demonstrating that all 150 L3 cells succeeded in tuning to contextually predictable features. Furthermore, Figure 8B shows that cross-correlations between basal outputs of different L3 cells residing in the same mesocolumn are low, indicating that these cells tuned to diverse canonical features.

**Figure 7.**
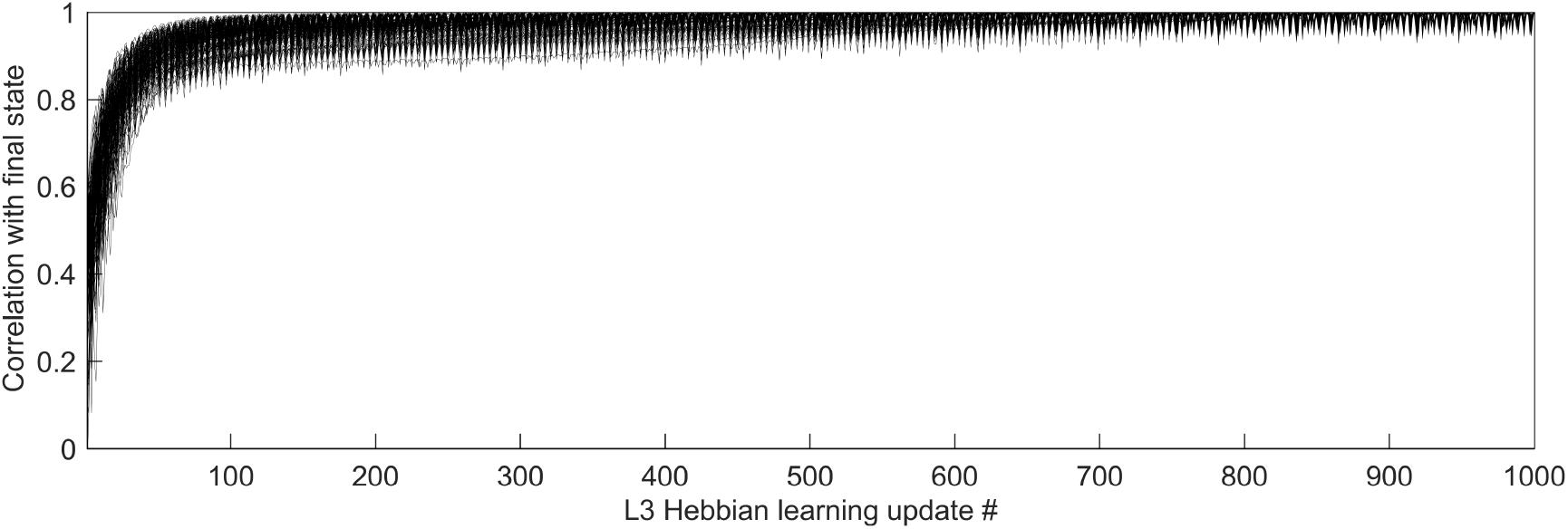
Time-course of L3 cells’ development of feature selectivities. Hebbian connections of the basal and apical dendrites of L3 cells were developed by modifying them iteratively over 1000 steps based on pre- and post-synaptic activity correlations computed in response to 5000 visual input patterns. To see how quickly the cells converge to their final connectional patterns, responses of each cell to a particular set of test input patterns were obtained after completion of 1000 connection updates, and these responses were correlated with responses to the same test set obtained after each connection update prior to the final one, thus using the correlation coefficient to express the similarity of the cell’s tuning at each update step to its final tuning. The plot shows superimposed the progressions of these correlation coefficients of all 150 L3 cells, from the first update to the last, revealing fast convergence to the final state without any meandering around.

**Figure 8.**
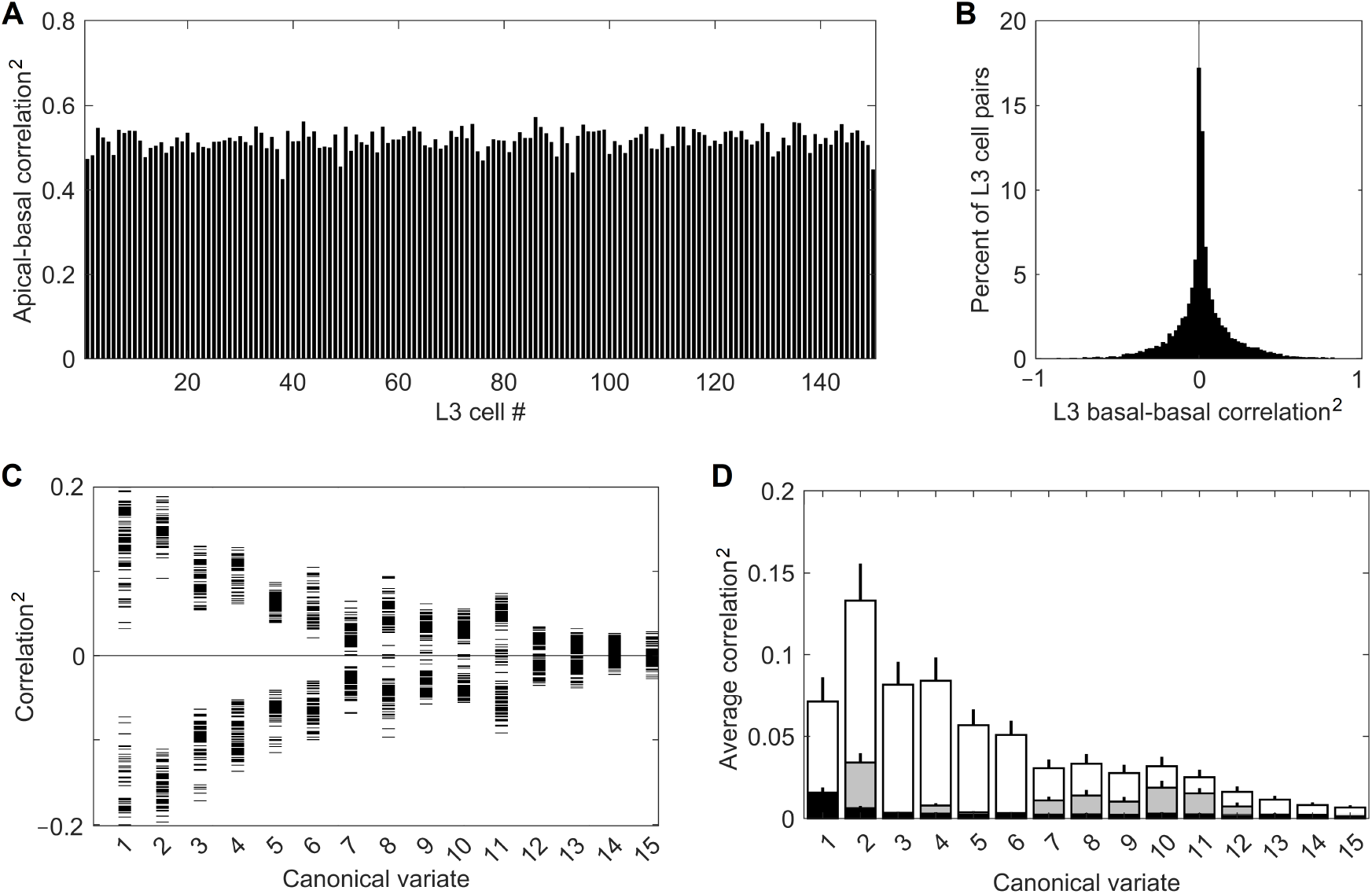
Successful tuning of L3 cells to contextually predictable features. **(A)** Uniformly high correlation (squared) of outputs of the apical and basal dendrites of the 150 cells in the mesocolumn’s L3 compartment. **(B)** Distribution of pairwise correlations (squared, keeping the sign) of outputs of basal dendrites of all 150 L3 cells, revealing their low similarity. **(C)** Basal dendrite correlations (squared, keeping the sign) of all L3 cells with each of the first 15 canonical variates (horizontal tick marks), showing that each L3 cell acquired gradually declining sensitivity to each of the first 11 variates. **(D)** Average magnitude of correlations (squared) of canonical variates with outputs of the 150 L4 cells in the mesocolumn (black bars) and outputs of basal dendrites of the 150 L3 cells (white bars), revealing complete insensitivity of L4 cells to canonical variates. Also plotted are the average correlations of canonical variates with 150 L3 cells in the no-context L3 model (gray bars). Development of feature selectivities of L3 cells was repeated 10 times, each time starting by assigning initial connection weights to L4 cells at random and training them on a different randomly selected sequence of image patches, then extracting canonical variates using another randomly selected set of image patches, followed by the same in L3. The bars in the plot show the means and SEM of the average correlations determined in the 10 runs. In all runs, L3 cells developed similar feature selectivities.

Figure 8C provides some insight into the nature of the features chosen by the 150 L3 cells. It plots correlations of the basal dendrite of each L3 cell in the central mesocolumn with each of the first 15 canonical variates computed for that mesocolumn. The plot shows that each L3 cell developed either positive or negative sensitivity to each of the first 11 canonical variates, declining gradually from the first to the last variate. Combined with information in Figure 8B, this indicates that L3 cells picked different mixes of positive and negative sensitivities to the 11 variates. If we view the canonical feature space defined by the first 11 variates as an 11-dimensional hypercube, Figure 8BC indicates that L3 cells picked different corners of this hypercube. Unlike L3 cells, L4 cells do not have any preferential sensitivity to the canonical variates (compare white- and black-shaded bars in Figure 8D). The prominent preferential sensitivity of L3 cells is the product of L3 self-organization. As our no-context L3 model shows, L3 self-organization without contextual guidance from surrounding columns also can to some degree enhance cells’ sensitivity to canonical variates (compare black- and gray-shaded bars in Figure 8D), but much less than under contextual guidance.

When evaluated by their responses to moving images of sinewave gratings, 80% of L3 cells fall in the complex-cell category, whereas 20% of L3 cells fall in the simple-cell category (Figure 9A). This suggests that as a group, L3 cells in a mesocolumn should be able to represent both the orientation and position of grating images in their output activity vector 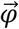 (Equation 4). To show how well they can do it, we expose the RF of the central mesocolumn to a grating pattern of randomly selected orientation, spatial frequency, and position in the RF, and compute the 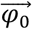 response it evokes in the 150 L3 cells in the central mesocolumn. We then rotate the grating pattern by a randomly chosen angle α, compute the new 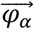 response of L3 cells, and measure the angle between the two L3 output vectors 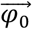 and 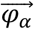. Figure 9B plots the average L3 angle as a function of the angle between the two gratings (gray curve). For a comparison, we also plot the average angle computed for the output vectors of the 150 L4 cells in the central mesocolumn (black curve). In Figure 9C, instead of rotating, we translate the grating pattern by a randomly chosen fraction (phase) of the grating’s period, compute the new response of L4 and L3 cells, and measure the angle between the two L3 output vectors and between the two L4 output vectors. Figure 9C plots the average L3 (gray) and L4 (black) angles as a function of the phase shift between the two gratings. The plots show that both L4 and L3 output vectors can discriminate even small differences in gratings’ orientation or position in the RF. It is interesting to note that even at the maximal orientation (90°) or spatial phase (180°) differences between two gratings, L3 output vectors show less than maximal (90°) separation, reflecting the fact that other than for their orientation or phase, the two gratings are the same.

**Figure 9.**
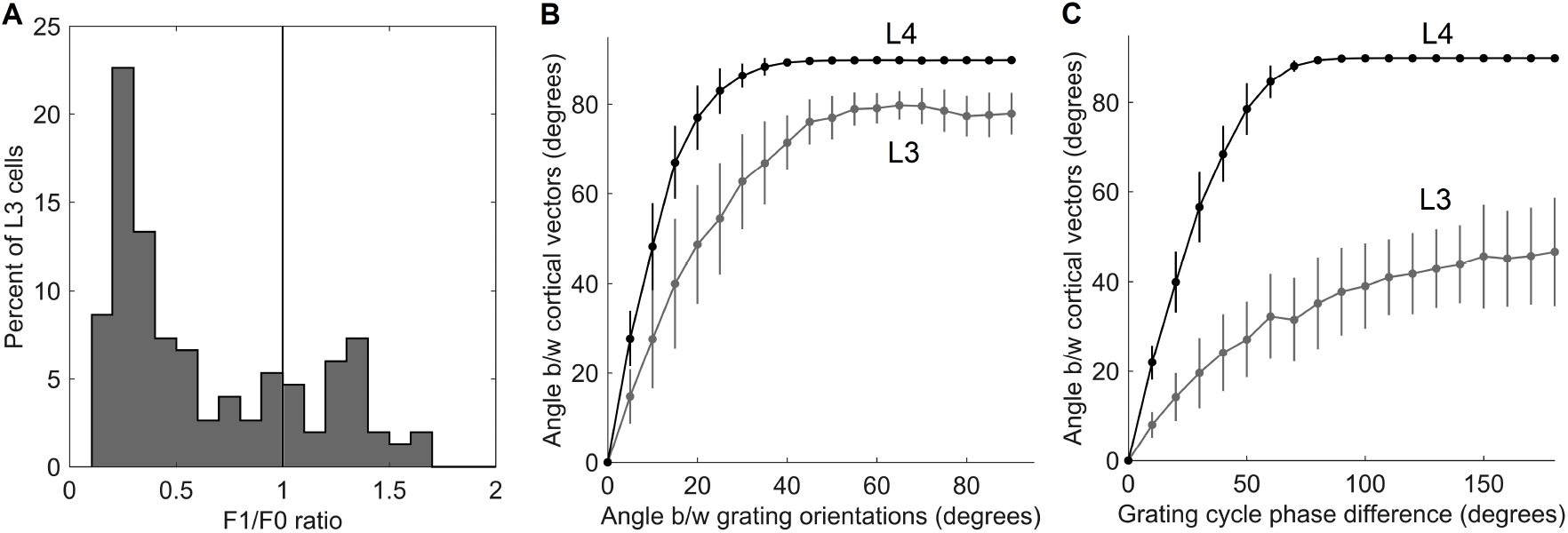
Responses of L3 cells to grating patterns. **(A)** F1/F0 scores of the 150 cells in the mesocolumn’s L3 compartment, spanning the range from clearly simple-cell (> 1) to clearly complex-cell (<1) categories. This distribution of ratios resembles that found in the upper layers in the real V1 cortex (Figure 1A, Kim and Freeman, 2016). **(B)** Discrimination of the grating orientations by the output vector of the 150 cells in the mesocolumn’s L4 compartment (black curve) and the 150 cells in the mesocolumn’s L3 compartment (gray curve). The average angle between two L4 or L3 output vectors is plotted as a function of the angle between orientations of the two compared gratings. **(C)** Discrimination of the grating placements by the output vector of the 150 cells in the mesocolumn’s L4 compartment (black curve) and the 150 cells in the mesocolumn’s L3 compartment (gray curve). The average angle between two L4 or L3 output vectors is plotted as a function of the phase shift of the two compared gratings. Vertical bars are standard deviations.

Figure 10 demonstrates the importance of contextual guidance for the development of biologically realistic feature properties in L3 cells. To test orientation tuning of L4 and L3 cells in the model, each cell was stimulated with moving grating patterns of the optimal spatial frequency and the full 180° range of orientations. The tightness of the cell’s orientation tuning was expressed by the standard half-width and half-height (HWHH) of the orientation tuning curve. Figure 10 plots the F1/F0 ratio determined for each cell against its HWHH. The model’s L4 cells are shown as blue circles, L3 cells as red dots, and L3 cells of the no-context model as green dots, revealing that all L4 cells are most tuned to orientation (average HWHH = 18°, matching real cat V1) and belong to the simple-cell category, L3 cells are also well-tuned to orientation and have biologically accurate proportion of simple- and complex cell categories, whereas L3 cells in the no-context model fail do develop translational invariance and have poor orientation tuning.

**Figure 10.**
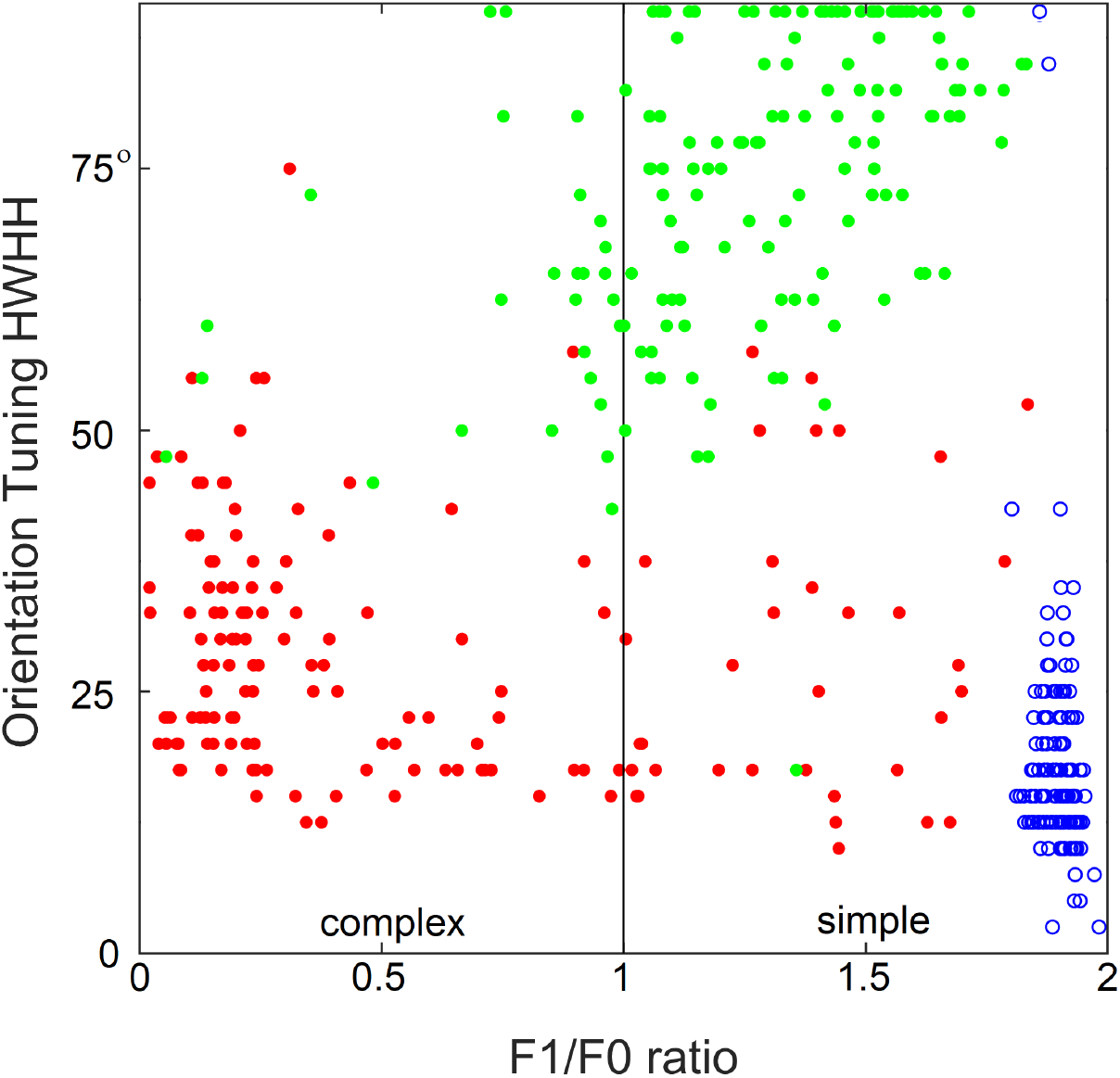
Importance of contextual guidance for the development of biologically realistic feature properties in L3 cells. Plotted against each other are the F1/F0 ratio and half-width at half-height (HWHH) of orientation tuning of the model’s L4 cells (blue circles) and contextually guided L3 cells (red dots), as well as L3 cells of the no-context model (green dots), showing that L4 cells and contextually guided L3 cells acquire biologically correct orientation tuning and translational invariance properties, whereas in the absence of contextual guidance L3 cells fail to do so.

#### 3.2.3 L3 emergent properties

In the real cortex, long-range horizontal connections link cortical columns separated by up to several millimeters in a cortical area. They preferentially link cortical sites that share similar functional properties but have non-overlapping RFs (Gilbert and Wiesel, 1983; DeFelipe *et al*., 1986; Lund *et al*., 1993; Burton and Fabri, 1995; Bosking et al., 1997). The fact that input patterns encountered by mesocolumns in their RFs possess contextually predictable features makes it possible for such long-range horizontal connections to establish Hebbian links between distant cortical columns even though they have non-overlapping RFs. When L3 cells tune to contextually predictable features, they become correlated with similar L3 cells in surrounding columns in their stimulation-evoked activities. Figure 11 shows the magnitude of such correlations between L3 cells in the central mesocolumn and functionally identical L3 cells in the first and second ring of surrounding mesocolumns. For a comparison, Figure 11 also shows that even in the first ring of mesocolumns, functionally identical L4 cells have very low correlations, which means that they would not be able to establish lateral Hebbian connections.

**Figure 11.**
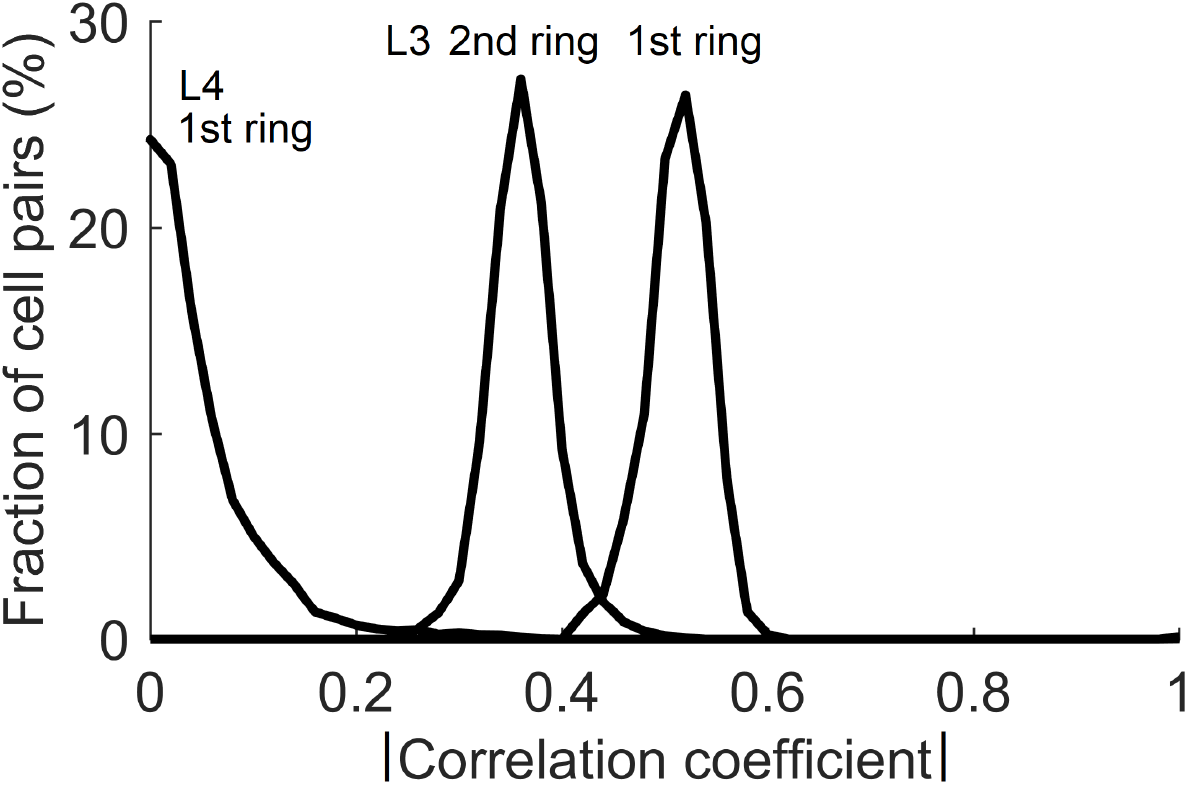
Correlations of cells in the central and surrounding macrocolumns in response to natural images. Plotted are distributions of magnitudes of correlation between functionally identical L4 and L3 cells in the central vs. the 1^st^ and 2^nd^ rings of surrounding macrocolumns. Substantial correlations at the level of L3 offer a substrate for growing Hebbian short- and long-range horizontal connections.

Canonical features learned by L3 cells characterize input patterns in many different ways that reflect the orderly aspects of the sensed outside world. This makes it possible for different input patterns, which are in some way objectively related, to be preferentially clustered in the L3 output space. We demonstrate this clustering tendency on an example of 4 different 512×512 pixel texture images shown in Figure 12A. We exposed the field of 19 mesocolumns to 500 randomly picked locations in each of the 4 texture images, and for each location we averaged the responses of L3 cells tuned to the same feature across 19 mesocolumns, resulting in a 150-dimensional feature vector representation of the imaged texture field. We next did Principal Component Analysis (PCA) on the 500×4 feature vectors. In Figure 12B, we plot the computed scores of the first 3 principal components, color-coding them according to the texture images from which they originated. This 3-D plot reveals that L3 responses to the 4 different textures occupy non-overlapping regions in the principal components space. For a comparison, Figure 12C shows the same plot for responses of thalamic LGN cells, which provided the input to the 19 cortical mesocolumns. As expected, LGN responses to the 4 different textures show no sign of preferential clustering.

**Figure 12.**
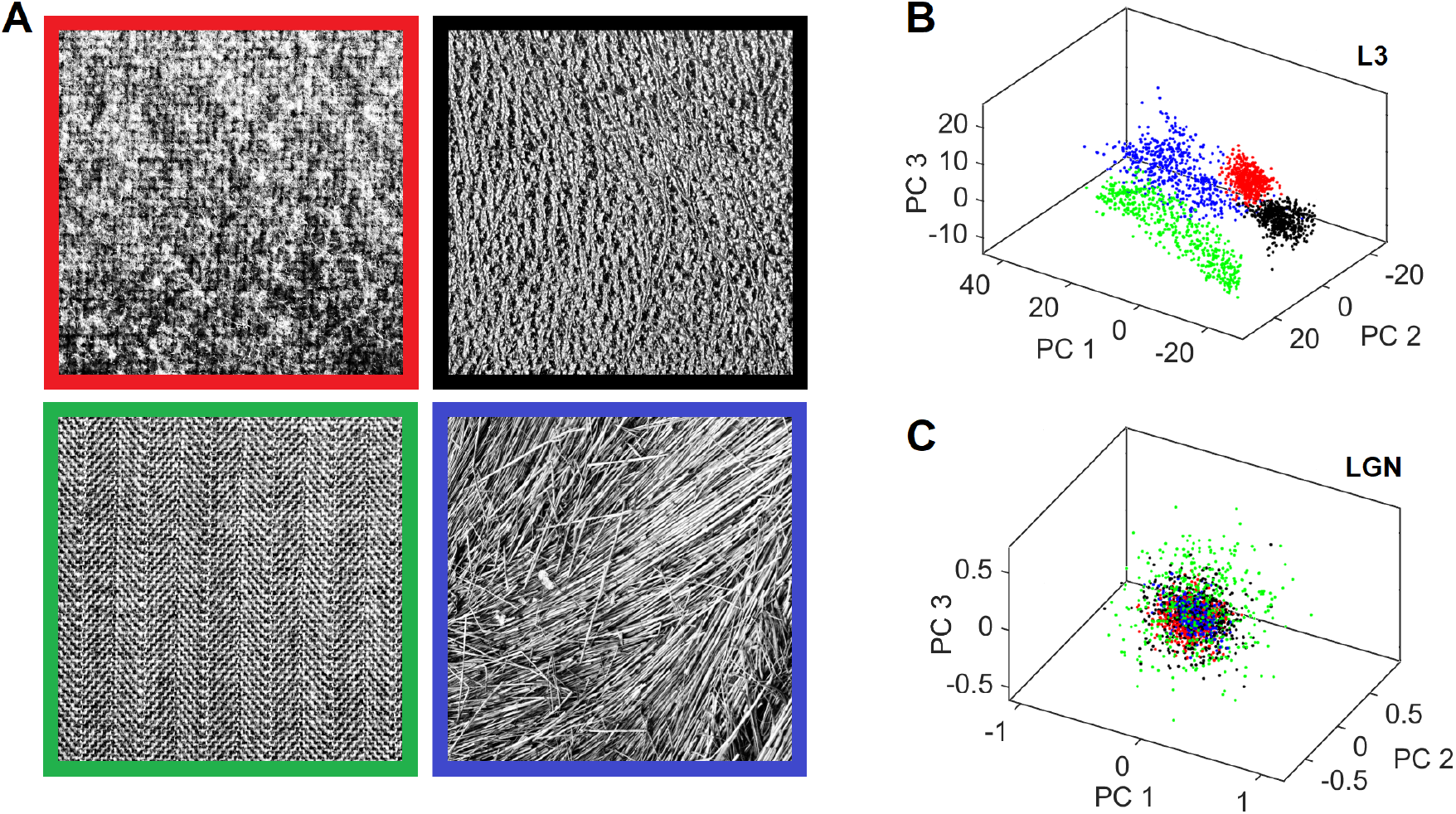
Separate clustering of different textures in the L3 output space but not in the L4 input space. **(A)** Four texture images used for demonstration. The viewing window of the central and 2 rings of macrocolumns (Figure 3B) was placed at 500 random locations in each of these images to obtain responses of all their L3 cells, yielding a 150-dimensional activity vector for each location. These vectors were converted to principal component scores by performing PCA on the 2000 vectors. **(B)** 3D plot of the scores of the first 3 principal components, color-coding each L3 activity vector by the frame color of its source texture image. L3 vectors coming from the same texture image cluster separately from other vectors, reflecting prominent visual differences among the 4 textures. **(C)** 3D scores plot of the afferent input vectors from LGN layer to L4, revealing that these activity vectors are all mixed together.

We expressed the similarity of L3 output vectors evoked in response to different randomly picked locations in the same texture image by computing correlations between pairs of such vectors. Figure 13 plots average correlations (squared, keeping the sign) for each of the 13 texture images in the Brodatz (1966) database. Also plotted are average correlations computed for the L4 response vectors and vectors of 15 significant canonical variates. As the plot shows, unlike L4 vectors, both L3 cell and canonical variate vectors evoked by different views of the same texture show substantial similarity. Also noticeable is that L3 cells consistently show greater similarity than canonical variates (compare red and green bars), demonstrating the advantages of the overcomplete representation of the mesocolumn’s 15-D canonical feature space by 150 L3 cells.

**Figure 13.**
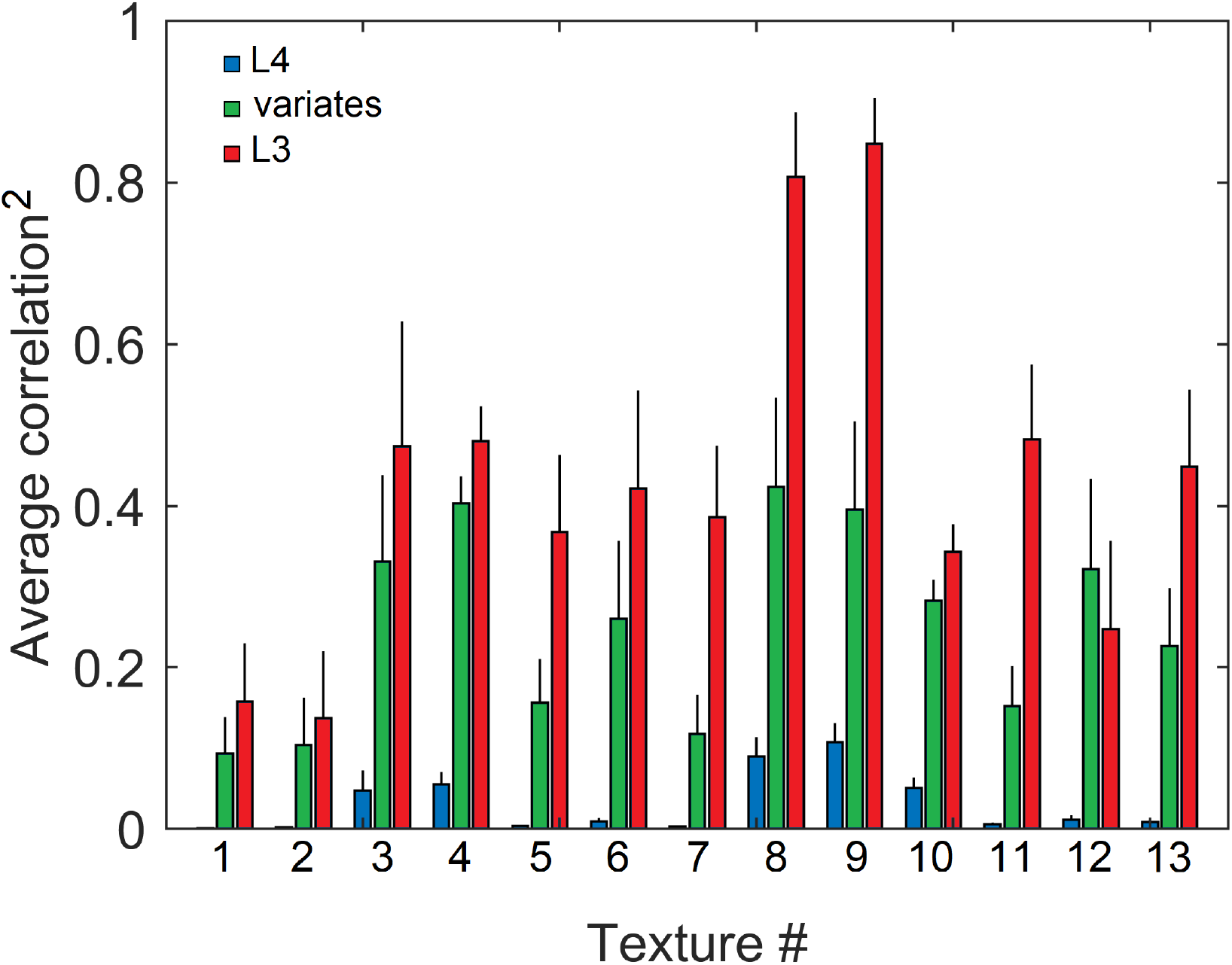
Similarity of output activity patterns evoked in L3, but not in L4, by different locations in the same texture image, reflecting their shared higher-order statistics. Responses of L4 and L3 cells in 19 mesocolumns, as well as their canonical variates, were obtained for 200 randomly placed locations in each of the 13 texture images in the Brodatz database, similar to Figure 12. Response profiles obtained from each texture image were cross-correlated and the average of these correlations (and standard deviation) were plotted separately for L4 (blue bars), canonical variates (green bars), and L3 (red bars) responses, showing that L4 responses had little, if any, similarity (the average correlation across the 13 images = 0.03), whereas L3 responses were more similar than canonical variate responses (13-image averages of 0.43 and 0.25, respectively).

## 4 DISCUSSION

### 4.1 Model accomplishments

The results of our cortical model simulations support biological plausibility of the proposed mechanism of cortical feature tuning and offer new insights into the nature of information extraction by neocortical networks. While potential usefulness of contextual guidance for feature tuning has long been recognized (see Introduction), so far it has only been explored at an abstract level or using greatly reduced “toy” models. In this paper we explore actual biological means by which neocortex tunes its neurons to contextually predictable features. Such means are likely to be the backbone of functional organization of cortical columns.

The starting point in our biologically grounded exploration was consideration that the tuned features have to be nonlinear. Our proposed two-stage solution is that, similar to artificial neural networks, feature tuning relies on hidden layer-like preprocessing, performed by the afferent input layer L4. That is, a local group of L4 neurons together perform a nonlinear transform of their thalamic inputs, which is akin to a basic RBF transform. Such transform accomplishes pluripotent function linearization, thus allowing L3 cells in the second stage to extract their features by simple linear summation of their L4 inputs.

Simplifying the task of feature tuning to that of linear operation over the L4 transform allowed us to determine – using an objective-function optimizing algorithm derived in Section 2.4 – that an RF of a representative V1 cortical column is likely to possess around 15 independent contextually predictable features, which we called canonical variates (Figure 4B). Furthermore, we determined that the transform-performing group of L4 cells will need at least 150 members in order to maximize the contextual predictability of all the canonical variates in its RF (Figure 6). Such a number is much greater than the 30-60 excitatory cells found in the L4 compartment of a single 0.05mm-diameter cortical minicolumn (which is the narrowest columnar entity in the neocortex), but much smaller than the total number of excitatory L4 cells in the 0.5mm-diameter macrocolumn. This finding leads us to propose a new class of cortical columns, the 0.15mm-diameter mesocolumn. It is estimated to comprise a local group of 7 minicolumns, all innervated by the same bundle of afferent axons, and to have 200-400 L4 excitatory cells, which together perform the pluripotent function linearizing transform of the mesocolumn’s afferent input.

Moving to the second stage of cortical feature extraction, which takes place in the upper layers, both anatomical evidence and model studies indicate that an L3 pyramidal cell extracts its feature from the L4 input it receives from not just its own mesocolumn, but from a macrocolumn-size group of 7 mesocolumns. Anatomical segregation of the ascending L4 inputs to the basal dendrites and the long-range horizontal inputs to the apical dendrites of L3 pyramidal cells, combined with their separate spike-generating synaptic input integrating centers allows basal and apical dendrites to have separate – classical and extra-classical – RFs and develop RF properties that will maximize covariance of the cell’s apical and basal outputs (Kording and Konig, 2000; Kay and Phillips, 2011). Our model simulations, based on Hebbian plasticity of apical and basal synapses, show that contextual inputs to the apical dendrites readily drive basal dendrites to select contextually predictable (i.e., canonical) features in their classical RFs. Similarly to real V1 cortex, 80% of model L3 cells acquire complex-cell RF properties while 20% acquire simple-cell properties (Gilbert, 1977; Kim and Freeman, 2016). Overall, the design of the model and its emergent properties are fully consistent with the known properties of cortical organization.

If cells in the mesocolumn’s L3 compartment did not push each other to select different features, they would all tune exclusively to the first – most predictable and thus most attractive – canonical variate. However, diversification pressures drive L3 cells to choose the second best solution. Rather than tuning to one or a mix of few of the canonical variates, all L3 cells in the mesocolumn become sensitive to all first 11 variates. This sensitivity declines gradually from the 1^st^ to the 11^th^ variate in all cells (Figure 8C). For each variate, all L3 cells develop approximately the same correlation with it but they differ in the sign of that correlation. Thus, each L3 cell carries maximal information it can about all first 11 variates (rather than emphasizing a subset of variates), with each variate contributing either positively or negatively in the pattern unique to that cell. As a result, L3 can be considered as approximating Hadamard-like domain transform of the first 11 canonical variates, decomposing them into a set of constituent functions (canonical features) over all variates. The L3 transform differs from Hadamard transform in that its transform functions are not orthogonal, and their number (200-400) greatly exceeds the number of variates (11). That is, L3 generates an overcomplete representation of canonical variates.

Considered geometrically, diversification pressures among cells in the mesocolumn’s L3 compartment drive them to choose different preferred directions in the mesocolumn’s canonical feature subspace. If we view the canonical feature space defined by the first 11 variates as an 11-dimensional hypercube, we find that L3 cells pick different corners of this hypercube. Such a hypercube will have 2048 corners to choose from. It is intriguing that while 200-400 L3 cells in a mesocolumn will not be able to pick all these corners, the larger columnar entity comprising a group of 7 mesocolumns – together making up a macrocolumn and sharing L4 input – will have just the right number of L3 cells for such a task.

As pointed out in Introduction, the orderly – as evidenced by their contextual predictability – nature of canonical features reflects the orderly structures in the environment. In tuning selectively to canonical features, L3 performs selective filtering of the information it receives from L4, emphasizing information about orderly aspects of the sensed environment and downplaying local, likely to be insignificant or distracting, information. Despite selective filtering and overcomplete representation of the canonical feature subspace, L3 output preserves excellent discrimination capabilities (Figure 9) while acquiring novel categorization/abstraction ability to preferentially cluster in the L3 output space different input patterns that are in some way objectively related (Figure 12). Furthermore, reduced sensitivity of L3 output to distracting irrelevant details should help the L4 in the next cortical area to minimize the Curse of Dimensionality and to succeed in the next round of pluripotent function linearization, and for the next L3 to find higher-order canonical features.

### 4.2 The model’s antecedents

The general idea of using spatiotemporal coherence to discover useful regularities in inputs was introduced by Becker and Hinton (1992) and later elaborated by Becker (1996). Their IMAX learning procedure discovers regularities in multiview inputs by maximizing mutual information between outputs of two nonlinear multilayer network modules that receive nonoverlapping, but spatially or temporally related, input samples, thus tuning to higher-order input features reflecting common distal causes in the external world. Details of IMAX design, however, make it unsuitable for implementation in the cerebral cortex (Becker, 1996). Phillips and Singer (1997) suggested a way of making computation of mutual information biologically more plausible, and it is one of the cornerstones of their Coherent Infomax theory. They consider abstract local processors, loosely analogous to unspecified local cortical circuits, that receive both the afferent input from their RFs and lateral (contextual field) input from other such local processors. The contextual field input guides local processors to tune to those stimulus features in their RFs that are predictably related to the context in which they occur. According to Coherent Infomax, contextual inputs can be used not only to guide learning but, importantly, also modulate short-term processing of sensory information. Phillips and Singer (1997) derived a particular mechanism for how contextually-guided learning might be accomplished. Unfortunately, that mechanism is limited in its practical utility due to its inability to search for nonlinear correlations. In its later development, Kay and Phillips (2011) showed that Coherent Infomax is consistent with a particular Bayesian interpretation for the contextual guidance of learning and processing and suggested learning rules that are more computationally feasible within systems composed of very many local processors.

Rather than invoking abstract local processors, Kording and Konig (2000) proposed that contextual guidance of feature tuning is implemented in individual pyramidal neurons, in which the apical dendrite acts – in addition to the soma – as a second site of integration capable of generating action potentials. Synaptic inputs to the soma site, coming from the cell’s RF, mainly determine the output activity of the post-synaptic neuron. Contextual inputs to the apical site gate synaptic plasticity. This separation makes it possible for contextual information to avoid confounding the effects of processing and learning. In “toy” simulations of such 2-site neurons receiving nonoverlapping but correlated inputs to their somata while sending their “teaching” outputs to each other’s apical site, cells learned to represent only the coherent part of the input, which would be expected to be relevant to the processing at higher stages. Kording and Konig termed their design Relevant Infomax.

To explain how 2-site pyramidal neurons might be able to tune to nonlinear features in their inputs, the challenge which was not addressed by the Kording and Konig model, Favorov and Ryder (2004) proposed that since dendritic trees are fundamentally nonlinear integrators, they might be able to operate functionally as error backpropagating multilayer perceptrons (MLP). In their SINBAD (acronym for *Set of INteracting BAckpropagating Dendrites*) neuron model (Ryder and Favorov, 2001), the apical dendrite in each pyramidal cell functions as one MLP and the basal dendrites function as the second MLP, using each other’s output activities as their reciprocal backpropagating teaching signals. While SINBAD cells are very powerful in discovering high-order nonlinear regularities hidden in multiview sensory inputs, effectively approximating Gebelein’s maximal correlation (Kursun and Favorov, 2010), it has become clear that they are not biologically feasible because, while action potentials do backpropagate from the initial axon segment up the apical dendrite, their experimentally observed amplitude modulation is not consistent with what would be required in the error signal. Furthermore, this design depends on a complete separation of the inputs to the apical and basal dendrites, which is not observed in the real cortex. Instead, a much more biologically appealing solution for the necessity of tuning cells to nonlinear features is to make use of pluripotent function linearization in L4 (Favorov and Kursun, 2011), followed by linear learning in L3, as is explored in this paper.

### 4.3 Model limitations

The model of contextual guidance of feature selection explored in this paper is not complete. In addition to spatial context, which was investigated here, contextual guidance can come from temporal context in which orderly features occur, as well as from higher-level understanding of the overall situation. In this paper, we only used static images and thus confined ourselves to spatial features of orderly structures, leaving temporal features of orderly processes for later studies. We anticipate that studies of feature acquisition under temporal contextual guidance and feedback from higher-level cortical areas will make it necessary to expand our current L4-L3 model by adding deep layers and layer 2, resulting in a cortical column model incorporating all cortical layers.

The biological realism of neurons modeled in this paper is not complete. Unlike real neurons, which have binary outputs and are either excitatory or inhibitory, but not both, the modeled cells have outputs that are continuous variables in a negative-positive range and have connections that in the process of learning can change their sign. Adding this degree of biological realism to the model will be insightful, but we do not expect it to negate the lessons learned using the current model. Also, some of the mathematical techniques used in the model, such as normalization of the connection weights in Equations 17, 23 and 25, might only approximate the true homeostatic mechanisms in the cortex (e.g., Turrigiano et al., 1998) and should be investigated further.

Sensory cortical columns are engaged not only in feature extraction and sensory information transmission to higher cortical areas, but also in other tasks, such as across-column binding by selective spike synchronization (Uhlhaas et al., 2009; Singer and Lazar, 2016), dynamic contrast enhancement and focused attention (Schummers et al., 2005; Tommerdahl et al., 2010; Tallon-Baudry, 2012), predictive computation (Bubic et al., 2010; Favorov et al., 2015; Marvan and Phillips, 2024; George et al., 2025), etc. Correspondingly, output of real pyramidal cells in L3 is determined not only by synaptic integration of L4 inputs by the basal dendrites, as was done in the current paper, but also by local excitatory and inhibitory inputs, input from the apical dendrite, and other sources (Angelucci and Bressloff, 2006). Our current model lacks all this machinery since its sole purpose was to investigate mechanisms determining classical RF and feature tuning properties of cortical neurons. However, assuming that our proposed mesocolumn-based mechanism of 2-stage feature extraction is biologically realistic, our current model provides a starting point, constraints, and guidance in building a progressively more comprehensive model of cortical functional organization.

## Funding

This work was supported, in part, by National Science Foundation grant 2435093 and by the J. Scott McFadyen Fund for Excellence in Parkinson’s Disease Research

## Acknowledgments

We thank Drs. Richard Murrow and Tim Challener for valuable discussions and comments on the manuscript.

## Notes

### Competing Interest Statement

The authors have declared no competing interest.

### Summary of Updates

This version of the manuscript has been substantially revised in response to reviewer feedback. We clarified that the canonical variates are constructed by maximizing correlations, and then showed how Hebbian learning distributes these compact dimensions into single-neuron selectivities, forming an overcomplete representation of contextual covariates. The experimental scope has been broadened to include a more diverse dataset. Additional analyses assess robustness to parameter choices, initialization, and convergence, demonstrating that the model's behavior is stable across configurations. Finally, comparisons with related models highlight the distinctiveness and advantages of our framework.

## References

Alpaydin E (2014). Introduction to machine learning, third edition. The MIT Press, Cambridge.

Angelucci A, Bressloff PC (2006) Contribution of feedforward, lateral and feedback connections to the classical receptive field center and extra-classical receptive field surround of primate V1 neurons. Prog. Brain Res. 154: 93–120.

Barlow HB (1992) The biological role of neocortex. In Information Processing in the Cortex, Aertsen A and Braitenberg V (eds), Springer, Berlin, pp. 53–80.

Beaulieu C, Colonnier M (1983) The number of neurons in the different laminae of the binocular and monocular regions of area 17 in the cat. J. Comp. Neurol. 217: 337–344.

Becker S (1996) Mutual Information Maximization: Models of Cortical Self-Organization. Network 7: 7–31.

Becker S, Hinton GE (1992). Self-organizing neural network that discovers surfaces in random-dot stereograms. Nature 355:161–163.

Bernander O, Koch C, Douglas RJ (1994) Amplification and linearization of distal synaptic input to cortical pyramidal cells J. Neurophysiol. 72: 2743–2753.

Bosking WH, Zhang Y, Schofield B, Fitzpatrick D (1997) Orientation selectivity and arrangement of horizontal connections in tree shrew striate cortex. J. Neurosci. 17:2112–2127.

Brodatz P (1966) Textures: A Photographic Album for Artists and Designers. Dover Publications.

Bruno RM, Simons DJ (2002) Feedforward mechanisms of excitatory and inhibitory cortical receptive fields. J. Neurosci. 22: 10966–10975.

Bubic A, von Cramon DY, Schubotz RI (2010) Prediction, cognition and the brain. Front. Hum. Neurosci. 4:25.

Budd JML (2000) Inhibitory basket cell synaptic input to layer IV simple cells in cat striate visual cortex (area 17): a quantitative analysis of connectivity. Vis. Neurosci. 17: 331–343.

Burton H, Fabri M (1995) Ipsilateral intracortical connections of physiologically defined cutaneous representations in areas 3b and 1 of macaque monkeys: projections in the vicinity of the central sulcus. J. Comp. Neurol. 355: 508–538.

Buxhoeveden DP, Casanova MF (2002) The minicolumn hypothesis in neuroscience. Brain 125:935–951.

Callaway EM (2004) Feedforward, feedback and inhibitory connections in primate visual cortex. Neural Networks 17: 625–632.

Cauller LJ, Connors BW (1994) Synaptic physiology of horizontal afferents to layer I in slices of rat SI neocortex J. Neurosci. 14: 751–762.

Cruikshank SJ, Lewis TJ, Connors BW (2007) Synaptic basis for intense thalamocortical activation of feedforward inhibitory cells in neocortex. Nat. Neurosci. 10: 462–468.

Da Costa NM, Martin KA (2010) Whose cortical column would that be? Front. Neuroanat. 4:16.

Deco G, Obradovic D. Decorrelated Hebbian learning for clustering and function approximation. Neural Comput. 7: 338–348, 1995.

DeFelipe J, Conley M, Jones EG (1986) Long-range focal collateralization of axons arising from corticocortical cells in monkey sensory-motor cortex. J. Neurosci. 6: 3749–3766.

DeFelipe J, Ballesteros-Yanez I, Inda MC, Munoz A (2006) Double-bouquet cells in the monkey and human cerebral cortex with special reference to areas 17 and 18. Prog. Brain Res. 154:15–32.

de Sa V, Ballard DH (1998) Category learning through multimodality sensing. Neural Comput. 10: 1097–1117.

DiCarlo JJ, Cox DD (2007) Untangling invariant object recognition. Trends Cogn. Sci. 11, 333–341.

Egger V, Feldmeyer D, Sakmann B (1999) Coincidence detection and changes of synaptic efficacy in spiny stellate neurons in rat barrel cortex. Nat. Neurosci. 2: 1098–1105.

Favorov OV, Diamond ME (1990) Demonstration of discrete place-defined columns – segregates - in the cat SI. J. Comp. Neurol. 298: 97–112.

Favorov OV, Kelly DG (1996a) Local receptive field diversity within cortical neuronal populations. In: Somesthesis and the Neurobiology of the Somatosensory Cortex. O. Franzen, R. Johansson and L. Terenius (eds.), Birkhauser Verlag AB, Basel, pp. 395–408.

Favorov OV, Kelly DG (1996b) Stimulus-response diversity in local neuronal populations of the cerebral cortex. NeuroReport 7: 2293–2301.

Favorov OV, Kursun O (2011) Neocortical layer 4 as a pluripotent function linearizer. J. Neurophysiol. 105:1342–1360.

Favorov OV, Whitsel BL, Tommerdahl M (2015) Discrete, place-defined macrocolumns in somatosensory cortex: lessons for modular organization of the cerebral cortex. In: Recent advances on the modular organization of the cortex. M. F. Casanova and I. Opris (eds.), Springer Science+Business Media Dordrecht, pp. 143–156.

Favorov OV, Nilaweera WU, Miasnikov AA, Beloozerova IN (2015) Activity of somatosensory-responsive neurons in high subdivisions of SI cortex during locomotion. J. Neurosci. 35:7763–7776.

Favorov OV, Ryder D (2004) SINBAD: a neocortical mechanism for discovering environmental variables and regularities hidden in sensory input. Biol. Cybernetics 90: 191–202.

Felleman DJ, Van Essen DC (1991) Distributed hierarchical processing in the primate cerebral cortex. Cereb. Cortex 1: 1–47.

George D, Lazaro-Gredilla M, Lehrach W, Dedieu A, Zhou G, Marino J (2025) A detailed theory of thalamic and cortical microcircuits for predictive visual inference. Sci. Adv. 11, eadr6698.

Gilbert CD (1977) Laminar differences in receptive filed properties of cells in cat primary visual cortex. J. Physiol. 268: 391–421.

Gilbert CD, Wiesel TN (1983) Clustered intrinsic connections in cat visual cortex. J. Neurosci. 3:1116–1133.

Golub GH, Van Loan CF (2013) Matrix Computations (4th ed.). Johns Hopkins University Press.

Grubinger M, Clough P, Müller H, Deselaers T (2006) The IAPR TC-12 benchmark: a new evaluation resource for visual information systems. Proceedings of the OntoImage 2006 Language Resources For Content-Based Image Retrieval. Genoa, Italy. Vol. 5, May 2006, p. 10. Available at: https://www.mathworks.com/help/deeplearning/ug/data-sets-for-deep-learning.html

Hardoon D, Szedmak S, Shawe-Taylor J (2004) Canonical correlation analysis: an overview with application to learning methods. Neural Comput. 16: 2639–2664.

Hawkins J, Blakeslee S (2004) On Intelligence. New York: Owl Books.

Hirsch JA, Martinez LM, Pillai C, Alonso JM, Wang Q, Sommer FT (2003) Functionally distinct inhibitory neurons at the first stage of visual cortical processing. Nat. Neurosci. 6: 1300–1308.

Hotelling H (1936) Relations between two sets of variates. Biometrika 28: 312–377.

Hubel DH, Wiesel TN (1962) Receptive fields, binocular interactions and functional architecture in the cat’s visual cortex. J. Physiol. 160: 106–154.

Kay JW, Phillips WA (2011) Coherent Infomax as a computational goal for neural systems. Bull. Math. Biol. 73: 344–372.

Kim T, Freeman RD (2016) Direction selectivity of neurons in visual cortex is non-linear and laminar dependent. Eur. J. Neurosci. 43: 1389–1399.

Kurková V (2003) Universal approximators. In: The Handbook of Brain Theory and Neural Networks (2nd ed.), ed. Arbib MA. Cambridge, MA: MIT Press, p. 1180–1183.

Kursun O, Favorov OV (2010) Feature selection and extraction using an unsupervised biologically-suggested approximation to Gebelein’s maximal correlation. International Journal of Pattern Recognition and Artificial Intelligence 24:337–358.

Kursun O, Patooghy A, Poursani P, Favorov OV (2024) CG-CNN: Self-supervised feature extraction through contextual guidance and transfer Learning. IEEE Access12: 155851–155866.

Kyriazi H, Carvell GE, Brumberg JC, Simons DJ (1996) Quantitative effects of GABA and bicuculline methiodide on receptive field properties of neurons in real and simulated whisker barrels. J. Neurophysiol. 75: 547–560, 1996.

Kisvardy ZF, Martin KAC, Freund TF, Magloczky Z, Whitteridge D, Somogyi P (1986) Synaptic targets of HRP-filled layer III pyramidal cells in the cat striate cortex. Exp. Brain Res. 64:541–52.

Kording KP, Konig P (2000) Learning with two sites of synaptic integration. Network: Computation in Neural Systems 11: 25–39.

Kursun O, Alpaydin E, Favorov OV (2011) Canonical correlation analysis using within-class coupling. Pattern Recognition Letters 32: 134–144.

Larkum ME, Waters J, Sakmann B, Helmchen F (2007) Dendritic spikes in apical dendrites of neocortical layer 2/3 pyramidal neurons. J. Neurosci. 27:8999–9008.

Larkum ME, Zhu JJ, Sakmann B (1999) A new cellular mechanism for coupling inputs arriving at different cortical layers Nature 398: 338–341.

Lin TY, Maire M, Belongie S, Bourdev L, Girshick R, Hays J, Perona P, Ramanan D, Zitnick CL, Dollar P (2015) Microsoft COCO: Common Objects in Context. arXiv:1405.0312v3 [cs.CV] 10.48550/arXiv.1405.0312

Lowe D (2003) Radial basis function networks. In: The Handbook of Brain Theory and Neural Networks (2nd ed.), ed. Arbib MA. Cambridge, MA: MIT Press, p. 937–127–940.

Lubke J, Roth A, Feldmeyer D, Sakmann B (2003) Morphometric analysis of the columnar innervation domain of neurons connecting layer 4 and layer 2/3 of juvenile rat barrel cortex. Cereb. Cortex 13: 1051–1063.

Lund JS, Yoshioka T, Levitt JB (1993) Comparison of intrinsic connectivity in different areas of macaque monkey cerebral cortex. Cereb. Cortex 3: 148–162.

Markram H, Muller E, Ramaswamy S, et al. (2015) Reconstruction and simulation of neocortical microcircuitry. Cell 163: 456–492.

Marvan T, Phillips WA (2024) Cellular mechanisms of cooperative context-sensitive predictive inference. Curr. Research Neurobiol. 6:100129.

MATLAB (2023) Version R2023a. The MathWorks, Inc. https://www.mathworks.com/

Meyer HS, Wimmer VC, Oberlaender M, de Kock CPJ, Sakmann B, Helmstaedter M (2010) Number and laminar distribution of neurons in a thalamocortical projection column of rat vibrissal cortex. Cereb. Cortex 20: 2277–2286.

Mountcastle VB (1978) An organizing principle for cerebral function: the unit module and the distributed system. In: The Mindful Brain, ed. Edelman GM, Mountcastle VB. Cambridge, MA: MIT Press, p. 7–50.

Mountcastle VB (1997) The columnar organization of the neocortex. Brain 120: 701–722.

Park J, Sandberg IW (1991) Universal approximation using radial-basis-function networks. Neural Comput. 3: 246–257.

Petreanu L, Mao T, Sternson SM, Svoboda K (2009) The subcellular organization of neocortical excitatory connections. Nature 457:1142–1146.

Phillips WA, Singer W (1997) In search of common foundations for cortical computation. Behav. Brain Sci. 20: 657–683.

Riesenhuber M, Poggio T (1999) Hierarchical models of object recognition in cortex. Nat. Neurosci. 2, 1019–1025.

Ringach DL, Shapley RM, Hawken MJ (2002) Orientation selectivity in macaque V1: Diversity and laminar dependence. J. Neurosci. 22: 5639–5651.

Ritter H (2003) Self-organizing feature maps. In: The Handbook of Brain Theory and Neural Networks (2nd ed.), ed. Arbib MA. Cambridge, MA: MIT Press, p. 1005–1010.

Rockland KS, Pandya DN (1979) Laminar origins and terminations of cortical connections of the occipital lobe in the rhesus monkey. Brain Res. 179: 3–20.

Ryder D (2004) SINBAD neurosemantics: a theory of mental representation. Mind & Language 19: 211–240.

Ryder D, Favorov OV (2001) The new associationism: a neural explanation for the predictive powers of cerebral cortex. Brain and Mind 2: 161–194.

Sáez I, Friedlander MJ (2009) Plasticity between neuronal pairs in layer 4 of visual cortex varies with synapse state. J. Neurosci. 29: 15286–15298, 2009.

Schölkopf B, Smola AJ (2002) Learning with Kernels. Cambridge, MA: MIT Press.

Schummers J, Sharma J, Sur M (2005) Bottom-up and top-down dynamics in visual cortex. Prog. Brain Res. 149: 65–81.

Silberberg G, Markram H (2007) Disynaptic inhibition between neocortical pyramidal cells mediated by Martinotti cells. Neuron 53: 735–746.

Singer W, Lazar A (2016) Does the cerebral cortex exploit high-dimensional, non-linear dynamics for information processing? Front. Comput. Neurosci. 10:99.

Schiller J, Schiller Y, Stuart G, Sakmann B (1997) Calcium action potentials restricted to distal apical dendrites of rat neocortical pyramidal neurons J. Physiol. 505: 605–616.

Skottun BC, DeValois RL, Grosof DH, Movshon JA, Albrecht DG, Bonds AB (1991) Classifying simple and complex cells on the basis of response modulation. Vision Res. 31: 1078–1086.

Somers DC, Nelson SB, Sur M (1995) An emergent model of orientation selectivity in cat visual cortical simple cells. J. Neurosci. 15: 5448–5465.

Stone JV (1996) Learning perceptually salient visual parameters using spatiotemporal smoothness constraints. Neural Comput. 8: 1463–1492.

Stuart GJ, Spruston N (1998) Determinants of voltage attenuation in neocortical pyramidal neuron dendrites J. Neurosci. 18: 3501–3510.

Swadlow HA (2003) Fast-spiking interneurons and feedforward inhibition in awake sensory neocortex. Cereb. Cortex 13: 25–32.

Sun QQ, Huguenard JR, Prince DA (2006) Barrel cortex microcircuits: thalamocortical feedforward inhibition in spiny stellate cells is mediated by a small number of fast-spiking interneurons. J. Neurosci. 26: 1219–1230.

Tallon-Baudry C (2012) On neural mechanisms subserving consciousness and attention. Front. Psychol. 2:397.

Tommerdahl M, Favorov OV, Whitsel BL, Nakhle B, Gonchar YA (1993) Minicolumnar activation patterns in cat and monkey SI cortex. Cereb. Cortex 3: 399–411.

Tommerdahl M, Favorov OV, Whitsel BL (2010) Dynamic representations of the somatosensory cortex. Neurosci. Biobehav. Rev. 34: 160–170.

Turrigiano GG, Leslie KR, Desai NS, Rutherford LC, Nelson SB (1998) Activity-dependent scaling of quantal amplitude in neocortical neurons. Nature 391:892–896.

Vinje WE, Gallant JL (2000) Sparse coding and decorrelation in primary visual cortex during natural vision. Science 287: 1273–1276.

Watanabe S (1960) Information theoretical analysis of multivariate correlation. IBM Journal of Research and Development 4: 66–82.

Uhlhaas PJ, Pipa G, Lima B, Melloni L, Neuenschwander S, Nikolic D, Singer W (2009) Neural synchrony in cortical networks: history, concept and current status. Front. Integrative Neurosci. 3, Art 17, 1–19.

